# Molecular mechanism of Mad2 conformational conversion promoted by the Mad2-interaction motif of Cdc20

**DOI:** 10.1101/2024.03.03.583158

**Authors:** Conny W.H. Yu, Elyse S. Fischer, Joe G. Greener, Jing Yang, Ziguo Zhang, Stefan M.V. Freund, David Barford

## Abstract

During mitosis, unattached kinetochores trigger the spindle assembly checkpoint by promoting assembly of the mitotic checkpoint complex, a heterotetramer comprising Mad2, Cdc20, BubR1 and Bub3. Critical to this process is the kinetochore-mediated catalysis of an intrinsically slow conformational conversion of Mad2 from an open (O-Mad2) inactive state to a closed (C-Mad2) active state bound to Cdc20. These Mad2 conformational changes involve substantial remodelling of the N-terminal β1 strand and C-terminal β7/β8 hairpin. *In vitro*, the Mad2- interaction motif (MIM) of Cdc20 (Cdc20^MIM^) triggers rapid conversion of O- to C-Mad2, effectively removing the kinetic barrier for MCC assembly. How Cdc20^MIM^ directly induces Mad2 conversion remains unclear. In this study we demonstrate that the Cdc20^MIM^-binding site is inaccessible in O-Mad2. Time-resolved NMR and molecular dynamics simulations show how Mad2 conversion involves sequential conformational changes of flexible structural elements in O-Mad2, orchestrated by Cdc20^MIM^. Conversion is initiated by the β7/β8 hairpin of O-Mad2 transiently unfolding to expose a nascent Cdc20^MIM^-binding site. Engagement of Cdc20^MIM^ to this site promotes release of the β1 strand. We propose that initial conformational changes of the β7/β8 hairpin allows binding of Cdc20^MIM^ to a transient intermediate state of Mad2, thereby lowering the kinetic barrier to Mad2 conversion.

## Introduction

During mitosis, accurate segregation of sister chromatids to daughter cells is critical to maintaining the faithful inheritance of genetic material and genomic stability. Segregation commences at the onset of anaphase when the cohesion between sister chromatids is released. This process is triggered only when all sister chromatid pairs have achieved successful bipolar attachment to the mitotic spindle, and tension is exerted at the microtubule-kinetochore attachment site. Unattached kinetochores activate the spindle assembly checkpoint (SAC) ^1^ to inhibit the anaphase-promoting complex/cyclosome (APC/C) and delay anaphase onset. SAC signalling is mediated by the SAC effector, the mitotic checkpoint complex (MCC), a heterotetrameric complex consisting of Mad2, Cdc20, BubR1 and Bub3 ^2,3^, that binds to and inhibits APC/C^Cdc20^ (Refs. ^4–6^), reviewed in ^7–11^.

MCC assembly requires the conversion of the metamorphic protein Mad2 from an inactive O- Mad2 to an active C-Mad2 state that has high affinity for Cdc20 ^12–19^ (Fig. 1a-c). This C- Mad2:Cdc20 complex spontaneously binds BubR1:Bub3 to generate the MCC ^3,20,21^. Previous structural studies on these two states of Mad2, that have distinct secondary structure topologies, are summarised in Supplementary Fig. 1. O-Mad2 and C-Mad2 share a central core consisting of a three-stranded antiparallel β sheet (β4-β6), three α helices (αA-αC) and a β hairpin (β2/β3) (cyan in Fig. 1a-c). Whereas the central core of Mad2 remains mainly unchanged following the O- to C-Mad2 conversion, the N- and C-terminal segments undergo major remodelling. In O- Mad2, the unstructured N-terminus is followed by the five-residue β1 strand, whereas the C-terminus consists of the β7/β8 hairpin and an unstructured tail, with the β7/β8 hairpin pairing with β6 of the core β sheet. Upon conversion to C-Mad2, the β7/β8 hairpin is displaced from β6 and repositioned, together with the C-terminal tail, to form the β8’/β8” hairpin that pairs with β5 on the opposite side of the β sheet. The β8’/β8” hairpin substitutes for β1 of O-Mad2 that is displaced to extend the N-terminus of αA in C-Mad2 (blue in Fig. 1a-c). This rearrangement allows Cdc20^MIM^ to pair with β6 of the central β sheet. Importantly, the reconfigured C-terminus of C-Mad2 entraps Cdc20^MIM^ through a ‘safety-belt’ structure connecting β6 with β8’ (Ref. ^15^) (red in Fig. 1a-c). This conformational change of Mad2 is a conserved feature of HORMA domain proteins ^8,22,23^ in which a HORMA-domain closure motif (here Cdc20^MIM^) is topologically linked within the closed state.

**Figure 1.**
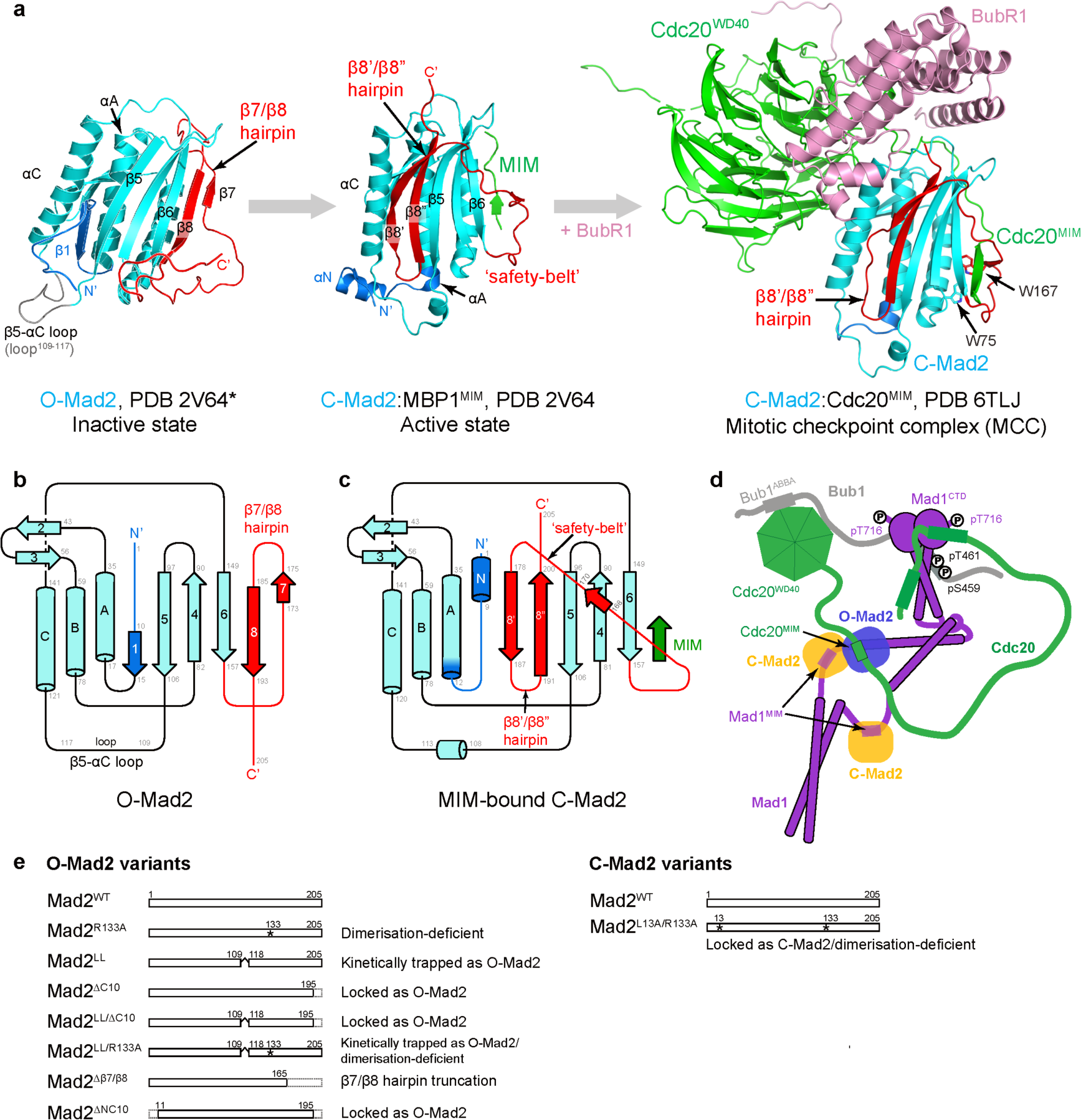
The conformational conversion of Mad2. a. Representative structural models of Mad2 in different conformations. The O-Mad2 structural model is based on the crystal structure of O-Mad2^LL^ (PDB 2V64, chain D) ^16^ and the missing segments were built using Modeller ^60^. The C-Mad2:MBP1^MIM^ structure is based on its crystal structure (PDB 2V64, chain A) ^16^. MBP1: Mad2-binding peptide1. The structure of C:Mad2:Cdc20^MIM^ in the BubR1:Cdc20:C-Mad2 complex is extracted from the cryo-EM structure of the APC/C:MCC complex (PDB 6TLJ) ^5,39^. The N-terminal residues 1-15 are in blue and the C-terminal residues 158-205 are in red to highlight the two segments that undergo significant conformational changes during the O- to C-Mad2 conversion. **b, c.** Secondary structure topology diagrams of O-Mad2 (**b**) and C-Mad2 (**c**). **d.** Schematic of the MCC-assembly scaffold. Mps1 kinase-phosphorylated Bub1 recruits the Mad1:C-Mad2 complex, O-Mad2 (blue) is recruited through self-dimerisation with C-Mad2 (orange), and Cdc20 interacts with Mps1-phosphorylated Mad1^CTD^ (C-terminal domain) and the ABBA motif of Bub1 ^31^. Through the formation of a tripartite Bub1:Mad1:Cdc20 complex, the MCC-assembly scaffold optimally positions Cdc20^MIM^ for its interaction with O-Mad2 and triggers the conversion of O-Mad2 to C-Mad2. **e.** Schematics of Mad2 variants used and discussed in this study. Mad2^R133A^, Mad2^LL^, Mad2^ΔC10^ and Mad2^LL/ΔC10^ were used for backbone assignment of O-Mad2.

The intrinsic rate of the O- to C-Mad2 conversion is extremely slow in solution (t_1/2_ of multiple hours) ^14,17,19,20^, whereas in cells, the SAC response is established within minutes ^24–26^.

Unattached kinetochores activate the SAC signal by catalysing formation of the MCC ^21^, in which the key catalysts of MCC-assembly are the protein kinase Mps1, the Mad1:C-Mad2 heterotetramer and Bub1 ^20,27,28^. These catalysts generate and contribute to an MCC-assembly scaffold that accelerates formation of the C-Mad2:Cdc20 complex ^27–32^. Mps1 activity, stimulated by the Ndc80 complex at unattached kinetochores ^33,34^, phosphorylates Knl1 and Bub1, thereby promoting Bub1 interactions with Mad1 and Cdc20. This scaffold both promotes the accessibility of Cdc20^MIM^ (through a proposed conformational change of Cdc20 ^28,30,35,36^) and juxtaposes Cdc20^MIM^ adjacent to its cryptic binding site on an O-Mad2 molecule interacting with C-Mad2 of the Mad1:C-Mad2 complex ^28,30–32^ (Fig. 1d). Accordingly, this catalyses formation of C-Mad2:Cdc20 ^28^. In support of this model, NMR experiments revealed that high concentrations of a peptide modelled on Cdc20^MIM^ (200 μM, in a two-fold access over O-Mad2) accelerated by at least 50-fold the entrapment of Cdc20^MIM^ by O-Mad2 to form C-Mad2:Cdc20^MIM^, as compared with the conversion rate of O-Mad2 to ‘empty’ C-Mad2 in the absence of MIM peptide ^31^.

Juxtaposing the accessible Cdc20^MIM^ adjacent to O-Mad2 by the MCC-assembly scaffold involves the dimerisation of an O-Mad2 molecule with Mad1-bound C-Mad2 ^28,30–32^. C-Mad2 binds to Cdc20^MIM^ and the MIM of Mad1 (Mad1^MIM^) through comparable mechanisms ^13–16^. *In vivo* studies demonstrated that disrupting the C-Mad2:O-Mad2 interaction abrogates SAC signalling ^16,18,37^. Additionally, the template model posits that C-Mad2:Mad1 contributes directly to catalysing MCC assembly by acting as a structural template for the conversion of O-Mad2 to C-Mad2 ^16,18^. This model is consistent with *in vitro* biochemical data showing that the O- to C- Mad2 conversion rate is modestly enhanced (3 to 4 fold) by dimerisation with C-Mad2 ^17,19^, and that purified chromosomes enhance formation of the MCC through a Mad2 template-dependent manner ^21^. However, *in vitro*, the rate enhancement contributed by O-Mad2:C-Mad2 dimerisation alone is substantially lower than the rate of MCC assembly in cells during an active SAC, consistent with observations that addition of the other MCC assembly catalysts accelerates MCC formation ^20,27,28^.

Despite progress in characterising Mad2 structurally, our understanding of the molecular mechanisms of the O- to C-Mad2 conversion, and how this is promoted by the MIM remains incomplete. In this study, we combined time-resolved NMR and molecular dynamics (MD) simulations to investigate the dynamics of structural elements in O-Mad2. We show that the Cdc20^MIM^-binding site on O-Mad2 is not accessible until exposed upon the transient unfolding of the flexible β7/β8 hairpin. By tracing the initial conversion rates of individual residues in O- Mad2, we demonstrate that Cdc20^MIM^ promotes the remodelling of the β1 strand, likely by facilitating its release from the core. Our data also indicate the existence of intermediate states during the O- to C-Mad2 conversion. We propose that conversion of O- to C-Mad2 in the presence of Cdc20^MIM^ is initiated by O-Mad2 transiently unfolding to expose the Cdc20^MIM^ binding site. This allows binding of Cdc20^MIM^ to an intermediate state of Mad2, stabilising this state, thereby lowering the kinetic barrier of O-Mad2 conversion to C-Mad2:Cdc20^MIM^.

## Results

### NMR fingerprinting of Mad2

We used time-resolved solution NMR to investigate the dynamic properties of the O- to C-Mad2 conversion. We collected ^1^H, ^15^N 2D HSQC spectra of Mad2 in different conformational states using different Mad2 variants (Fig. 1e), and in the presence of different MIM ligands. HSQC spectra are referred as protein ‘fingerprints’ since they are unique to different proteins. To gain structural insights on full-length Mad2, using the dimerisation-deficient mutant Mad2^R133A^ (Ref.^15^), we determined a near-complete assignment of Mad2 in its open state (O-Mad2), ‘empty’ closed state (C-Mad2) and ‘bound’ closed state (C-Mad2:Cdc20^MIM^). The assignments are described in Supplementary Fig. 2a-c and in Methods.

In the absence of MIM, O-Mad2 slowly converts to ‘empty’ C-Mad2 that structurally resembles the MIM-bound-C-Mad2 (Supplementary Fig. 3a). Despite their structural similarity, the NMR fingerprints for ‘empty’ C-Mad2 and the C-Mad2:Cdc20^MIM^ complex show large-scale chemical shift differences (Supplementary Fig. 2d) ^31^. These differences arise from the rearrangement of aromatic residues whose ring currents influence the nuclear environments of backbone amides in their close proximity (Fig. 2a). Likewise, despite the structural similarities of the ligand-bound C-Mad2 complexes, their 2D spectra are characteristic of specific MIM peptides (Supplementary Fig. 2e). Differences were observed for the indole NH resonances of Trp75 and Trp167 (Fig. 2b). In particular for Trp75, which is situated close to the MIM-binding site, significant perturbation was observed with bound ligand (Fig. 2a, b). The indole NH resonances from Mad2 tryptophan side chains were therefore used to identify the conformational state of the Mad2 MIM-binding site and its bound ligand.

**Figure 2.**
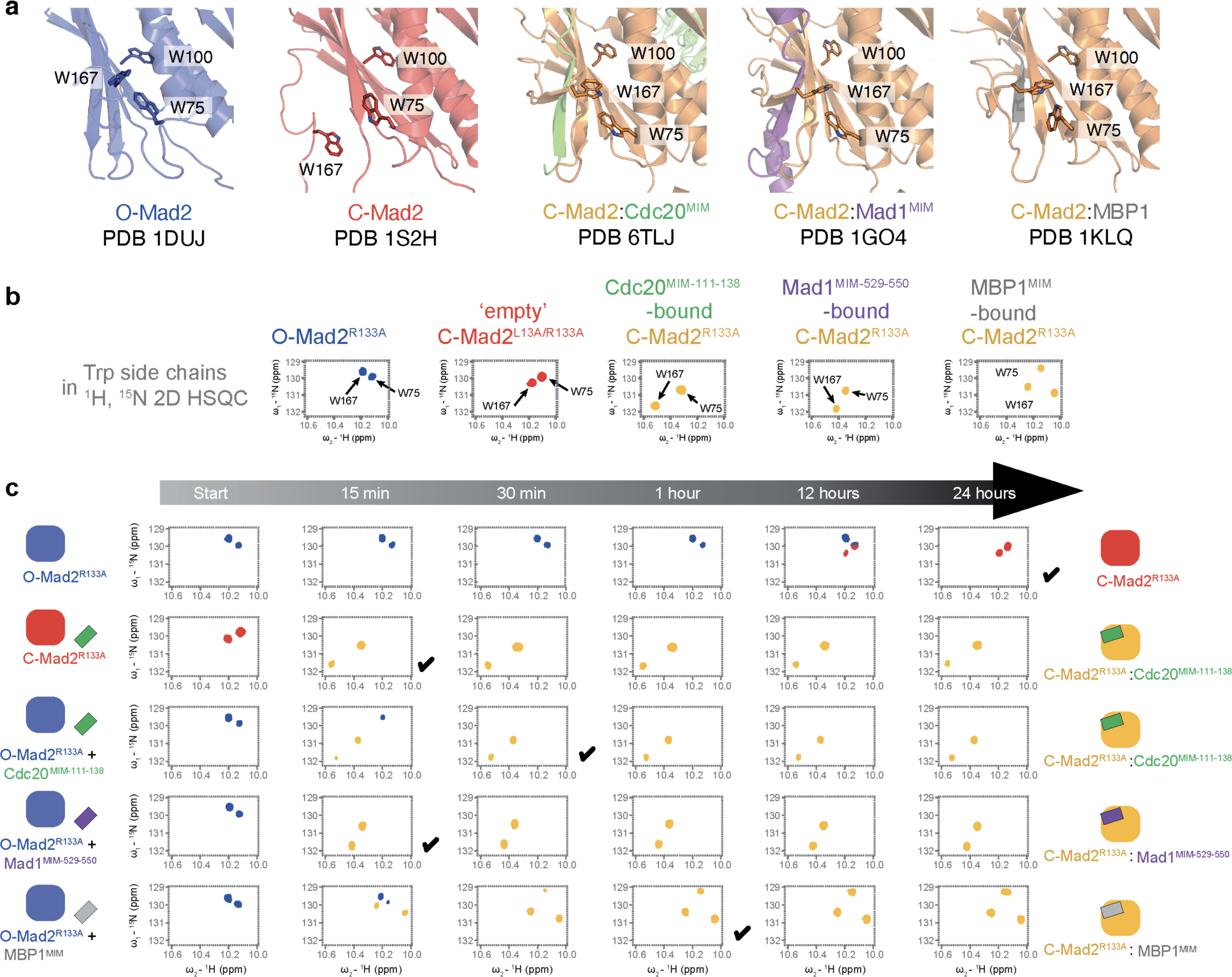
NMR fingerprinting of Mad2. a. Different structural orientations of Trp75 and Trp167 in Mad2 give distinct indole NH resonances in their NMR spectra. Trp167 is located at one end of the ‘safety-belt’ and is displaced during conversion. Trp75 is located on αB helix and is not involved in major conformational rearrangements. However, as Trp75 is situated near the MIM-binding site, significant perturbations were observed with different MIM ligands. **b.** Resonances from the indole NH of Trp75 and Trp167 have distinct chemical shifts in different conformations of Mad2. NMR spectra for O-Mad2^R133A^ and ‘empty’ C-Mad2^L13A/R133A^ and C-Mad2^R133A^:Cdc20^MIM-111-138^ are from ^31^ **c.** Mad2 conversion occurs at different rates in the presence of different MIM peptides. Using the side chain resonances of Trp75 and Trp167 as reporters, Mad2 conversion was traced in a 24 h time course at 25 °C. MIM peptides were added at two-fold molar excess to Mad2 (100 μM). Each spectrum required 12 min acquisition, therefore the 15 min time-points represent the earliest spectra in the time-resolved series. The schematics show the conformation of Mad2 as indicated by the tryptophan side-chain resonances. The full time-resolved spectra are shown in Supplementary Figs. 4 and 5. A black tick indicates when conversion is considered complete. NMR data for O-Mad2^R133A^ with and without Cdc20^MIM-111-138^ shown for comparison are from^31^.

We then investigated the secondary structures of the two Mad2 conformational states using their secondary chemical shifts. Secondary chemical shifts report deviations of the experimental Cα and Cβ chemical shifts from the theoretical values for a random coil. A positive value indicates helical propensity whereas a negative value indicates β-strand propensity ^38^. Overall, our secondary chemical shifts correlate well with the secondary structures of experimentally determined structures (Supplementary Fig. 3). For O-Mad2 we observed no secondary structure for the N- and C-terminal ten residues (Supplementary Fig. 3b). In solution, the N-terminal ten residues did not adopt a stable αN helix in either ‘empty’ C-Mad2 or C-Mad2:Cdc20^MIM^ (Supplementary Fig. 3c, d). A short αN helix observed in the crystal structure of C- Mad2:MBP1^MIM^ (MBP1^MIM^: MIM of Mad2-binding peptide1 ^13^) dimerised with O-Mad2 ^16^, is absent in the cryo-EM structure of the APC/C:MCC complex ^5,39^ or any other crystal structure of full-length C-Mad2 (Supplementary Fig. 1) ^40^.

### The MIM-binding site is inaccessible in O-Mad2

To understand whether Mad2 conversion rates are modulated by different C-Mad2-ligands, we used time-resolved NMR to determine the conversion rate of 100 µM O-Mad2^R133A^ at 25 °C. The indole NH resonances of Trp75 and Trp167 were used as reporters (Fig. 2b). We reported previously that in the presence of Cdc20^MIM-111–138^ (Cdc20^MIM^ peptide spanning Cdc20 residues 111-138), Mad2 conversion occurred in less than 30 min ^31^ (Fig. 2c and Supplementary Fig. 4a). In contrast, conversion took ∼24 h in the absence of Cdc20^MIM-111-138^ peptide ^31^ (Fig. 2c and Supplementary Fig. 4b). Thus, the MIM peptide accelerates the O- to C-Mad2 conversion by nearly 50-fold, comparable to the 35-fold acceleration of MCC assembly previously reported in the presence of full-length Cdc20 and the catalytic scaffold (Mad1:C-Mad2, Bub1:Bub3, Mps1)^28^. These data support the idea that an essential role of the MCC assembly catalysts is to present Cdc20^MIM^ at high effective concentration and optimal orientation proximal to the cryptic Cdc20^MIM^-binding site in O-Mad2. The O- to C-Mad2 conversion in the presence of Mad1^MIM-529-550^ (Mad1 residues 529-550) reached completion within 15 min (i.e. within the acquisition time of an individual HSQC spectrum), and 60 min in the presence of MBP1^MIM^, a phage-display MIM peptide selected for its enhanced C-Mad2 affinity ^13^ (Fig. 2c and Supplementary Fig. 5a, b). Our finding that Mad1^MIM-529-550^ induced the O- to C-Mad2 conversion agrees with an earlier NMR study showing that Mad1 residues 540-551 facilitate this conversion ^14^. We observed that ‘empty’ C-Mad2 rapidly adopted a ‘bound’ C-Mad2:Cdc20^MIM^ conformation on addition of Cdc20^MIM^ (Fig. 2c and Supplementary Fig. 4c), indicating that the MIM-binding site is easily accessible in ‘empty’ C-Mad2 for MIM peptides, consistent with prior observations that ‘empty’ C-Mad2 readily binds Cdc20^MIM^ when presented as short peptides ^17,28^. In contrast, ‘empty’ C- Mad2 does not readily associate with full-length Cdc20 ^28^. In ‘empty’ C-Mad2 the β7/β8 hairpin is displaced to expose the β6 strand for pairing with the MIM peptide ^14^ (Supplementary Fig. 3a). O-Mad2 that is unable to convert to C-Mad2 does not bind Cdc20^MIM^, as determined from NMR spectra when Cdc20^MIM^ was titrated into Mad2^ΔC10^ (Supplementary Figs. 6a-c and 7a), a Mad2 variant trapped in the open state ^18^, consistent with previous studies that Mad2^ΔC10^ cannot interact with full-length Cdc20 ^12,14,41^.

### An extended binding site for Cdc20^MIM^ involving linker residues 138-152

Cdc20^MIM-111-138^ had previously been used to study the O- to C-Mad2 conversion ^31^ (Fig. 2c and Supplementary Fig. 4a). However, it was unclear whether Cdc20-binding sites for Mad2 are limited to this region, given the extended linker connecting the MIM and WD40 domain of Cdc20 (residues 139-152) (Fig. 3a) that might interact with Mad2. Furthermore, residues N-terminal to Glu126 of Cdc20 were not resolved in the cryo-EM structure of the human APC/C:MCC complex ^5,39^, and it was unclear if these residues contribute to Mad2 binding and conversion. To identify potential additional Mad2-binding residues on Cdc20, and the corresponding Cdc20-binding sites on Mad2, we designed various length Cdc20^MIM^ peptides and compared their effects on Mad2-conversion rates by NMR (Fig. 3a, b and Supplementary Fig. 8). Cdc20^MIM^ spanning residues 123-137 (Cdc20^MIM-123–137^) triggered Mad2 conversion at a similar rate as Cdc20^MIM-111-138^ (Fig. 3b and Supplementary Figs. 4a and 8a). Moreover, there were no major chemical shift differences between the spectra of C-Mad2:Cdc20^MIM-123-137^ and C- Mad2:Cdc20^MIM-111-138^ (Supplementary Fig. 8b), indicating that residues 111-122 are not involved in either inducing the O- to C-Mad2 conversion or interacting with C-Mad2. In contrast, Cdc20^MIM-123-152^, that incorporates the PEG motif (residues 138-144), was more efficient at triggering Mad2 conversion and completely converted to C-Mad2 within 15 min (Fig. 3b and Supplementary Fig. 8c). This is in agreement with the finding that ablation of the PEG motif reduces the catalytic incorporation of Cdc20 into the MCC ^28^. However, Cdc20 residues 138-152 failed to induce Mad2 conversion after two hours, and therefore alone are insufficient to bind Mad2 (Fig. 3b and Supplementary Fig. 8d).

**Figure 3.**
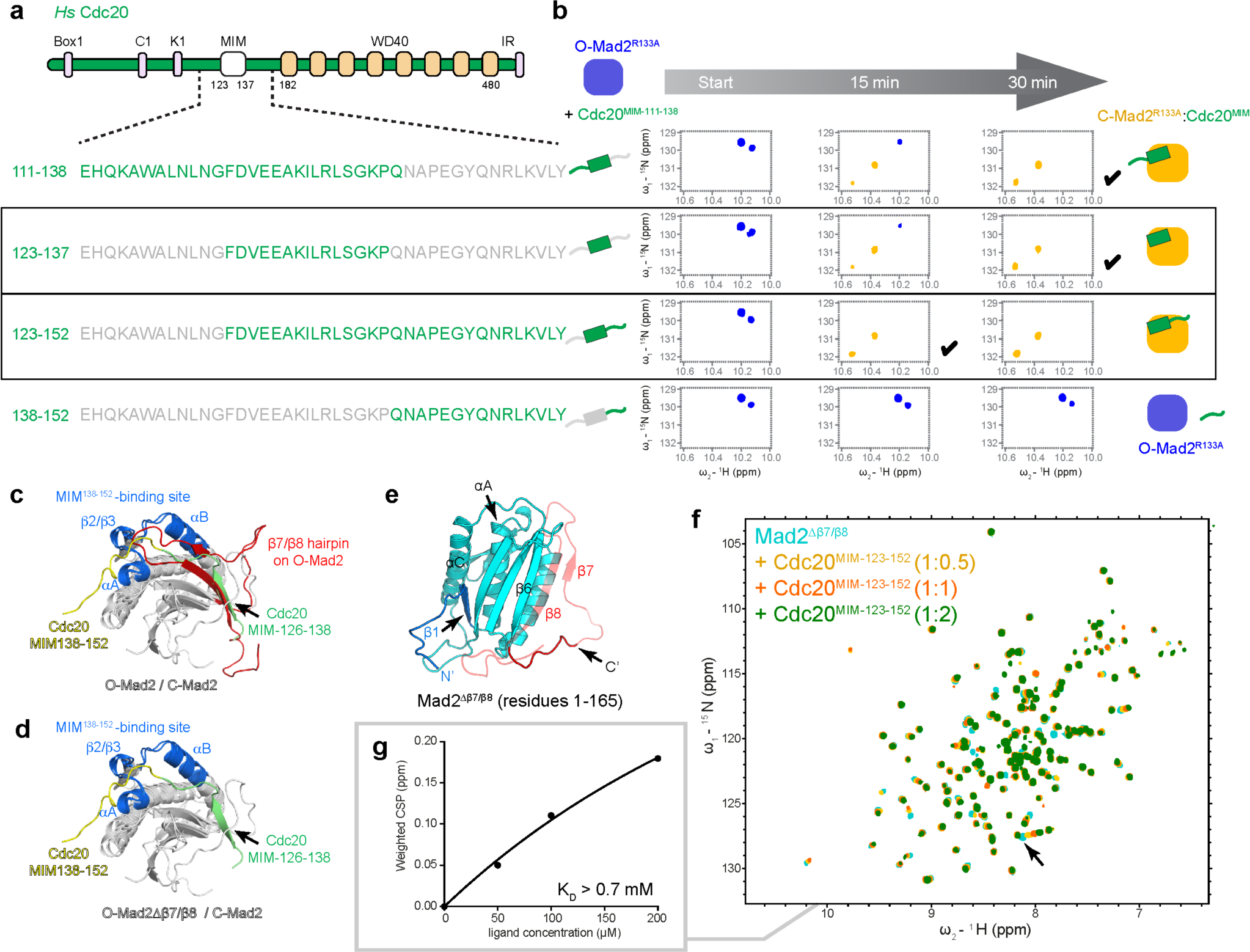
Mad2 conversion rates with different Cdc20^MIM^ peptides. a. Schematic of human Cdc20. Box1: Box1 motif; C1: C-box motif; K1: KEN box motif; MIM: Mad2-interacting motif, residues 123- 137; WD40: WD40 domain, residues 182-480; IR: IR tail. **b.** Mad2 conversion in the presence of different Cdc20^MIM^ peptides. Mad2 conversion was traced at 25 °C and the changes in the first 30 min are shown. The schematics illustrate the lengths of different MIM peptides, and the full spectra are shown in Supplementary Figs 4a and 8. A black tick indicates where conversion is considered complete. NMR data for O-Mad2^R133A^ with Cdc20^MIM-111-138^ shown for comparison are from ^31^. **c.** The Cdc20^MIM-138-152^-binding site was identified by comparing the spectra of C-Mad2:Cdc20^MIM-123-137^ with C-Mad2:Cdc20^MIM-123-152^ (boxed in **b**) and analysing their CSPs (Supplementary Figs. 8e, f). The Cdc20^MIM-138-152^-binding site (blue) is mapped onto the structure of C-Mad2:Cdc20 from the cryo-EM structure of the APC/C:MCC complex (PDB 6TLJ) ^5,39^. The O-Mad2 structural model is superimposed onto the C-Mad2 structure for comparison. Cdc20 residues 138-152 are in yellow and residues 126-138 are in green. The β7/β8 hairpin in O-Mad2 (red) blocks both the binding sites for Cdc20 residues 126-138 and residues 138-152. **d.** As in **c** except the β7/β8 hairpin in O-Mad2 is removed to expose the Cdc20 binding sites. **e.** Model of the Mad2^Δβ7/β8^ variant (residues 1-165; Λι166-205) to test Cdc20^MIM^-binding. The truncated residues 166-205 are shown with transparency. **f.** Cdc20^MIM-123-152^ was titrated into Mad2^Δβ7/β8^ and its binding was analysed by NMR. **g.** Resonance with the largest CSP in **f** (marked by an arrow) was used to estimate the binding affinity.

Comparing the spectra of C-Mad2:Cdc20^MIM-123-152^ and C-Mad2:Cdc20^MIM-123-137^ (Supplementary Fig. 8e) identified an additional Cdc20-binding site on Mad2 as revealed by chemical shift perturbations (CSPs) dependent on the additional Cdc20 residues 138-152 (Fig. 3c and Supplementary Fig. 8f). This Cdc20^MIM-138-152^-binding site comprises residues of the αA-β2 and β3-αB loops of Mad2. In the cryo-EM map of the APC/C:MCC complex ^5,39^, residues 134- 143 of Cdc20 are located in close proximity to this site. However, the resolution in this region of the cryo-EM map limited our analysis of the interaction between Cdc20^MIM-138-152^ and C-Mad2. The relatively small CSPs and lack of structural information on Cdc20^MIM-138-152^ suggested that residues 138-152 of Cdc20 only weakly associated with C-Mad2. Similar to the major MIM- binding site involving β6, the putative Cdc20^MIM-138-152^-binding site is blocked by the β7/β8 hairpin (coloured red in Fig. 3c). Overall, these results indicate that Cdc20 residues 123-138 constitute the core of Cdc20^MIM^, consistent with the APC/C:MCC cryo-EM structure ^5,39^.

We reasoned that MIM binding to Mad2 is blocked until the β7/β8 hairpin is displaced (Fig. 3c, d). To test this hypothesis, we generated a Mad2 variant with a deleted β7/β8 hairpin (Mad2^Δβ7/β8^: residues 1-165) (Fig. 3e). Mad2^Δβ7/β8^ adopted a stable globular fold, and due to the absence of the C-terminal residues, was unable to adopt a closed conformation (Fig. 3f).

Chemical shift perturbations induced by addition of Cdc20^MIM-123-152^ indicated a weak interaction between Cdc20^MIM-123-152^ and Mad2^Δβ7/β8^, with an estimated K_D_ of > 0.7 mM (Fig. 3f, g). This is in agreement with a previous study that showed a weak interaction between Mad2^Δβ7/β8^ and MBP1^MIM^ (Ref. ^40^). Our data indicate that the MIM-binding sites on O-Mad2 are blocked by the β7/β8 hairpin and that MIM binding requires the displacement of the β7/β8 hairpin.

### Unfolding the β7/β8 hairpin exposes the MIM-binding site

How Cdc20^MIM^ induces the O- to C-Mad2 conversion when the Cdc20-binding sites are not accessible in O-Mad2 was unclear. Due to the dynamic properties of O-Mad2, we hypothesised that MIM-binding sites might be transiently exposed on displacement of the β7/β8 hairpin. To assess this, we analysed the backbone dynamics of O-Mad2 using a combination of NMR relaxation experiments and MD simulations (Fig. 4a). NMR ^15^N transverse relaxation (T2) data provide information on backbone mobility on the picosecond to nanosecond timescale and are therefore a reliable diagnostic of protein dynamics. We measured the ^15^N T2 relaxation times for O-Mad2^ΔC10^ but were unable to collect equivalent data for O-Mad2^R133A^ because this variant slowly converted to C-Mad2 during data acquisition. The HSQC spectrum for O-Mad2^ΔC10^ overlaps with O-Mad2^WT^ and O-Mad2^R133A^ (Supplementary Fig. 6a). Assignment of the backbone amide resonances in O-Mad2 was less complete for the β7/β8 hairpin region (residues 160-190) due to their resonances suffering from exchange broadening, indicative of multiple conformational states. To fill these gaps, we performed MD simulations on both full-length O- Mad2 and O-Mad2^ΔC10^. A total of 5.4 µs of simulations were performed at 300 K. We then analysed the root mean square fluctuation of Cα atoms (Cα RMSF). Cα RMSF is a measure of the magnitude each Cα atom moves over the simulation and corresponds to residue flexibility (Fig. 4a). RMSD plots are shown in Supplementary Fig. 9. Due to the timescale of the O- to C- Mad2 conversion (∼24 h at 25 °C, Fig. 2d), these simulations could not model the entire open-to-closed conversion. The Cα RMSF from MD simulations of full-length O-Mad2 and O-Mad2^ΔC10^ are highly comparable (Fig. 4a), suggesting the backbone dynamics of O-Mad2 were not significantly disrupted by the C-terminal deletion. Importantly, MD Cα RMSF data and NMR T2 relaxation times were closely correlated (Fig. 4a). We therefore reasoned that the Cα RMSFs in our MD simulations reliably reflect the protein backbone dynamics of O-Mad2.

**Figure 4.**
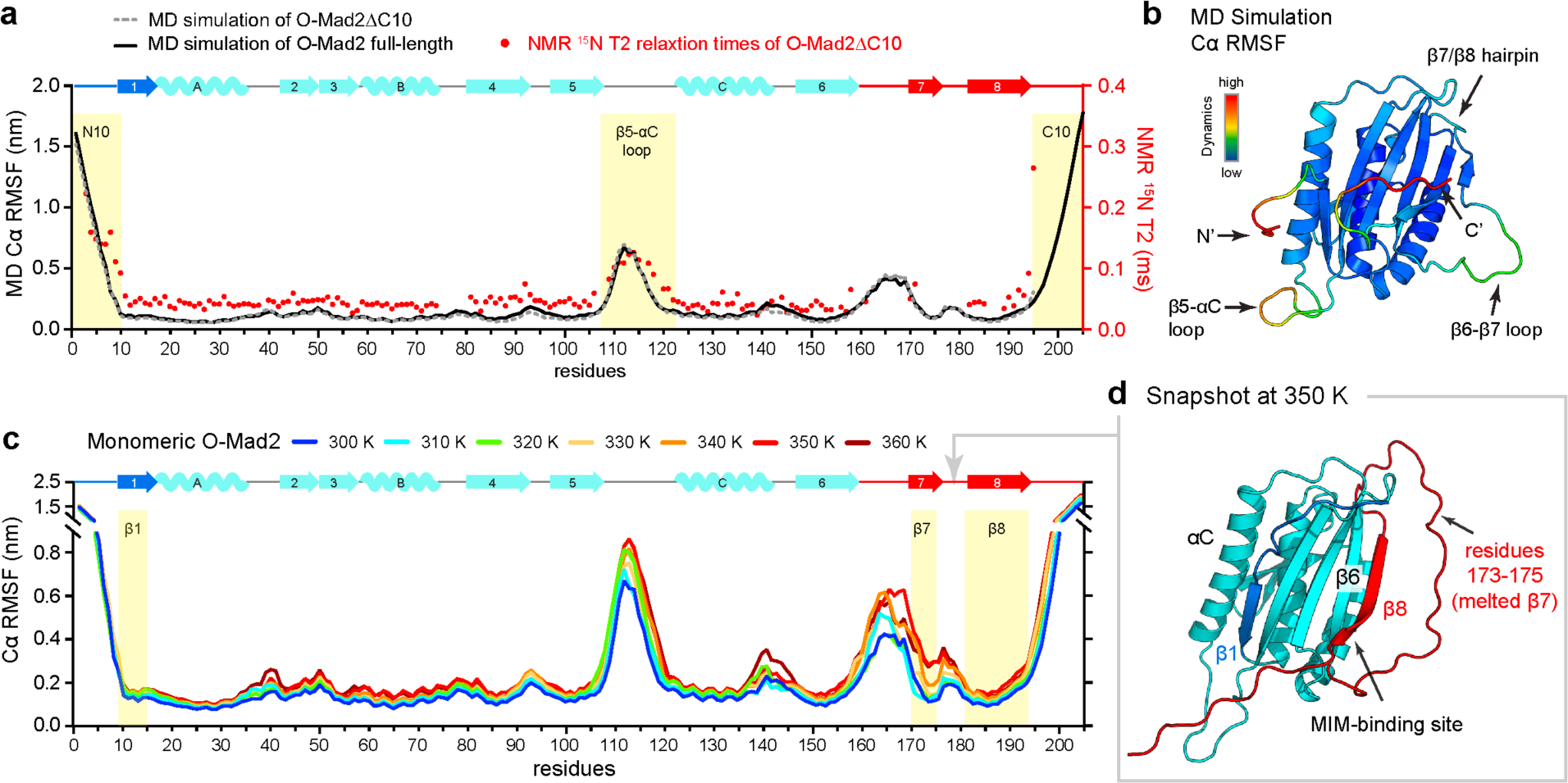
The C-terminal β7/β8 hairpin may transiently unfold to expose the MIM-binding site. NMR and MD simulations were used to study the dynamics of O-Mad2. A total of 5.4 µs of simulations were carried out for each variant, using six independent simulations of 0.9 µs. RMSD plots for all MD simulations are shown in Supplementary Fig. 9. **a.** ^15^N T2 relaxation times (red circles) were measured at 298 K and Cα RMSF were determined from MD simulations at 300 K for full-length O-Mad2 (black lines) and O-Mad2^ΔC10^ (dotted grey lines). Regions that are highly dynamic are highlighted. **b.** Dynamics data from MD simulations were mapped onto O-Mad2, with red being the most dynamic and blue being the least dynamic. **c.** Cα RMSF determined from MD simulations at 300-360 K for monomeric O-Mad2. **d.** A snapshot of the O-Mad2 simulation at 350 K acquired after 0.6 µs of simulation.

Our dynamics data indicated that O-Mad2 consists of a stable protein core with flexible regions comprising the ten residues at both N- and C-termini, the β5-αC loop and residues 160-185 (β6/β7 loop, β7 strand and β7/β8 loop) (Fig. 4a, b). A dynamic N-terminus (residues 1-10) agrees with our observations that there was no residual helical propensity for αN (Supplementary Fig. 3b). There is a sharp drop in backbone dynamics at residue 11 of the β1 strand (residues 11-15), suggesting that the β1 strand is stably bound to the core of O-Mad2. MD analysis indicated a highly flexible β6/β7 loop connecting the β6 strand with the β7/β8 hairpin. There is an overall increase in dynamics in this region, although with notable dips at the β7 and β8 strands (Fig. 4a). There are no T2 relaxation NMR data reporting on the dynamics of the C-terminal ten residues of O-Mad2, however the Cα RMSFs indicate a highly dynamic C-terminus, including residues 191-200 that remodel to become the β8” strand in C-Mad2 (Fig. 1b, c).

Upon the O- to C-Mad2 conversion, two structural elements are released from the core of O- Mad2: the N-terminal β1 strand, and the C-terminal β7/β8 hairpin (Fig. 1a). Both are directly connected to highly flexible termini (Fig. 4a), and we hypothesised that either element might be transiently displaced to initiate conversion. We reasoned that the melting of the short β7 strand (residues 173-175) is likely the first event in Mad2 conversion as β7 is held in place by only three main-chain hydrogen bonds. To test this hypothesis, we performed MD simulations at increasing temperatures to investigate which structural elements are least stable. We performed MD simulations on full-length O-Mad2, using a temperature range from 300 to 360 K (Fig. 4c and Supplementary Fig. 9). A total of 5.4 µs of simulations were performed at each temperature. The protein core of O-Mad2 remained stable from 300 to 340 K, consistent with the experimentally determined melting temperature of 342 K ^42^. At 350 K, β7 was simulated to dissociate from the core, resulting in an increased Cα RMSF (Fig. 4c, d). The C-terminal half of β8, which pairs with β6, was also found to be dynamic in all simulations, occasionally peeling off from β6 at 350 K (Fig. 4d). This would expose the binding site for MIM on the C-terminal half of β6. Meanwhile, β1 and the N-terminal half of β8 remained bound to the core even at the high simulated temperature of 360 K (Fig. 4c). The transient unfolding of the C-terminal β7/β8 hairpin would allow O-Mad2 to bind Cdc20^MIM^, however as Cdc20^MIM^ only weakly associates with Mad2^Δβ7/β8^ (Fig. 3f, g), the transient exposure of the MIM-binding site alone is likely not sufficient to secure Cdc20^MIM^ binding. A complete displacement of the β7/β8 hairpin, together with the β1 strand, is necessary to entrap Cdc20^MIM^ through the formation of the ‘safety-belt’. A flexible β7/β8 hairpin contrasting with a more rigid β1 strand is consistent with prior hydrogen deuterium exchange data revealing fast amide proton exchange for β8 and slow exchange for β1 ^43^. Finally, the C-terminus of the αC helix became more flexible at 360 K.

### Dynamic β5-αC loop of O-Mad2 modulates the O- to C-Mad2 conversion

To understand the contributions of the dynamic β5-αC loop in the O- to C-Mad2 conversion, we examined the loopless Mad2 mutant (Mad2^LL^) in which the β5-αC loop is truncated ^16^. This truncation creates a kinetic barrier to the remodelling of the β1 strand, severely reducing rates of conversion ^16^. We observed no binding of Cdc20^MIM-111-138^ to Mad2^LL^ in NMR spectra collected early in the time-course (Supplementary Fig. 6d). However, we noticed a slow time-dependent change in the NMR spectra indicating that O-Mad2^LL^ underwent conformational transition to C- Mad2^LL^ (Fig. 5 and Supplementary Fig. 7b), with the final spectrum of C-Mad2^LL^:Cdc20^MIM^ overlapping closely with that of full-length C-Mad2:Cdc20^MIM^ (Supplementary Fig. 7c, d). In the presence of Cdc20^MIM-111-138^, conversion of O-Mad2^LL^ to C-Mad2^LL^:Cdc20^MIM^ took ∼250 min, 8 times longer than the conversion of O-Mad2^R133A^ with Cdc20^MIM-111-138^ (Fig. 5). Given that the slow conversion of O-Mad2^LL^ to C-Mad2^LL^:Cdc20^MIM^ is caused by the shorter β5-αC loop posing a kinetic barrier to β1 remodelling ^16^, we conclude that Mad2 conversion is dependent on β1 remodelling.

**Figure 5.**
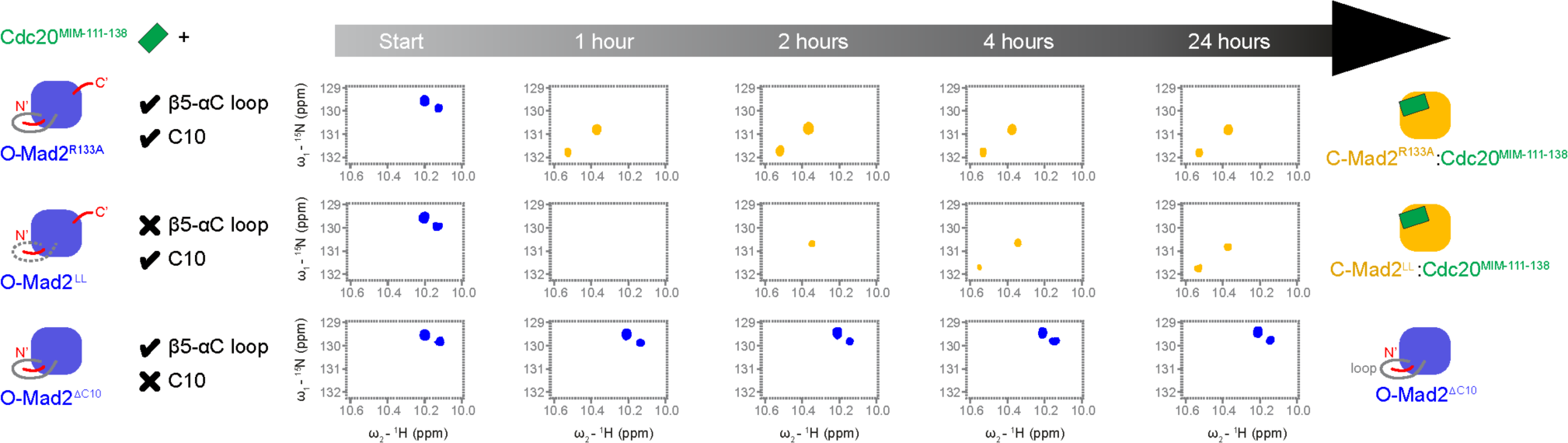
Dynamic β5-αC loop and C-terminal residues in O-Mad2 modulate Mad2 conversion. The O- to C-Mad2 conversion is slowed when the highly dynamic β5-αC loop is truncated (O-Mad2^LL^) and eliminated in the absence of the C-terminal ten residues (O-Mad2^ΔC10^). Full spectra for O-Mad2^LL^ and O- Mad2^βC10^ shown in Supplementary Fig. 7. Using the side chain resonances of Trp75 and Trp167 as reporters, Mad2 conversion was traced in a 24 h time course at 25 °C. Cdc20^MIM-111-138^ peptides were added at two-fold molar excess to Mad2 (100 μM). Severe line broadening was observed during conversion of O-Mad2^LL^, most likely due to dimerisation of O-Mad2^LL^ with the newly converted C- Mad2^LL^. NMR data for O-Mad2^R133A^ with Cdc20^MIM-111-138^ shown for comparison are from ^31^.

### O-Mad2 retains its dynamic and structural properties in the O-Mad2:C-Mad2 dimer

In the physiological context of MCC assembly, O-Mad2 dimerises with a Mad1-bound C-Mad2 ^14,17,19,40^ (Fig. 1d). We therefore investigated whether O-Mad2 undergoes a similar conversion to C-Mad2, when dimerised with C-Mad2, as we observed for the monomeric O-Mad2^R133A^ mutant. We first collected HSQC spectra of O-Mad2^WT^ in its monomeric state (Fig. 6a) and then monitored the changes in the spectra upon addition of Cdc20^MIM-123-152^ (Fig. 6b). O-Mad2^WT^ adopted a similar global fold to O-Mad2^R133A^, although with significant CSPs mapping to the β1 strand (Supplementary Fig. 10a, b), and converted to C-Mad2 upon binding Cdc20^MIM^ (Fig. 6b). The resultant C-Mad2^WT^:Cdc20^MIM^ generated comparable spectra to the dimerisation-deficient C-Mad2^L13A/R133A^:Cdc20^MIM^ (Fig. 6b). Size-exclusion chromatography coupled with multi-angle light scattering (SEC-MALS) data indicated the sample contained a mixture of monomeric and dimeric Mad2 (Fig. 6b inset), consistent with the increased NMR linewidths resulting from the higher molecular mass of the dimer. Because Cdc20^MIM^ peptide was added in excess and no O- Mad2 resonances were observed, we reasoned that the dimer detected in this experiment is a C- Mad2:C-Mad2 dimer, similar to that observed by others ^14,40^.

**Figure 6.**
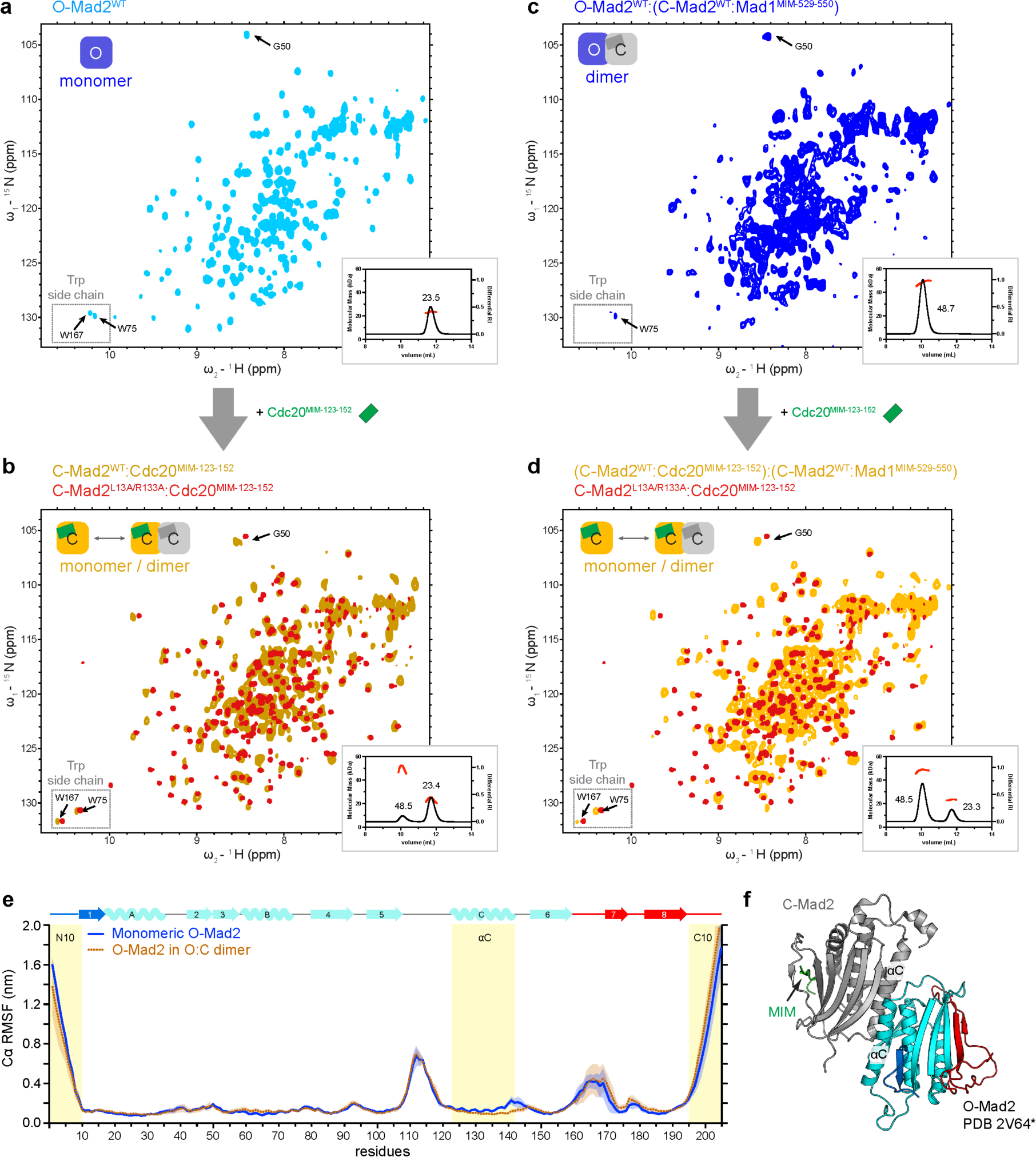
O-Mad2 retains its dynamic and structural properties in the O-Mad2:C-Mad2 dimer. a, c. HSQC spectra of 300 µM wild-type O-Mad2^WT^ (**a**, light blue) and O-Mad2^WT^:(C-Mad2^WT^:Mad1^MIM^) (**c**, blue). The schematics show the conformation of Mad2 as indicated by the tryptophan side-chain resonances and their oligomeric states as determined by SEC-MALS (insets). The SEC peaks are labelled with their molecular masses (kDa), where the theoretical molecular masses for monomeric and dimeric Mad2 are 23.5 and 47.0 kDa respectively. For the dimeric samples, only one of the monomers was isotopically labelled and observable by NMR. Distinct chemical shift differences were also observed for Gly50 in O-Mad2 and C-Mad2, and its resonances are marked in the spectra. Cdc20^MIM-123-152^ was added at two-fold molar excess to O-Mad2^WT^ (**a**), and O-Mad2^WT^:(C-Mad2^WT^:Mad1^MIM^) (**c**). **b, d.** This generated C-Mad2^WT^-Cdc20^MIM^ (**b**, brown), and (C-Mad2^WT^:Cdc20^MIM^):(C-Mad2^WT^:Mad1^MIM^) (**d**, yellow), respectively. Their spectra are overlaid with that of C-Mad2^L13A/R133A^:Cdc20^MIM^ (red) for comparison. **e.** Cα RMSF were determined from MD simulations at 300 K for monomeric O-Mad2 (blue line) and O-Mad2 in a O-Mad2:C-Mad2:Mad1^MIM^ dimer (brown dotted line). The continuous error bands indicate one standard deviation derived from six independent simulations. **f.** Model of O-Mad2:C- Mad2:Mad1^MIM^ based on the O-Mad2^LL^:C-Mad2:MBP1^MIM^ structure (PDB 2V64) ^16^ highlighting the αC helices at the dimeric interface.

We then investigated the O- to C- conversion of O-Mad2^WT^ in the context of the O-Mad2^WT^:C- Mad2^WT^:Mad1^MIM^ dimer. O-Mad2:C-Mad2:Mad1^MIM^ was prepared by mixing isotopically labelled O-Mad2^WT^ with an excess of unlabelled C-Mad2^WT^ bound to Mad1^MIM^ (Fig. 6c). This dimeric species was confirmed by SEC-MALS (Fig. 6c inset). The NMR signature of the Trp side chains of O-Mad2^WT^ in the context of O-Mad2:C-Mad2:Mad1^MIM^ corresponded closely with that of monomeric O-Mad2^WT^ (Supplementary Fig. 10c), indicating O-Mad2^WT^ adopted a similar global fold in both monomeric and dimeric states, and consistent with the overall similarity of the solution structure of O-Mad2^ΔNC10^ and O-Mad2 in the O-Mad2^LL^:C-Mad2:MBP1^MIM^ crystal structure (Supplementary Fig. 10e) ^12,16^. Severe line broadening was observed in the spectra of O-Mad2^WT^:C-Mad2:Mad1^MIM^, except for the flexible N-terminus (N10), β5-αC loop and β7/β8 hairpin (Supplementary Fig. 10d). This is a combined effect of the increased molecular mass of the dimer and conformational changes caused by dimerisation. The widespread line broadening across the sequence is comparable to the extensive chemical shift perturbations in O-Mad2 on dimerising with C-Mad2 ^44,45^. Similar to monomeric O-Mad2^WT^, O-Mad2^WT^ in the O-Mad2^WT^:C- Mad2:Mad1^MIM^ dimer converted to C-Mad2 upon binding Cdc20^MIM^ within 15 min (Fig. 6d).

The resulting C-Mad2:Cdc20^MIM^ adopted a similar fold to monomeric C- Mad2^L13A/R133A^:Cdc20^MIM^ (Fig. 6d), but the increased linewidths indicated the sample contained a mixture of monomeric and dimeric C-Mad2, as confirmed by SEC-MALS analysis (Fig. 6d inset).

To assess whether O-Mad2 dynamics are altered in the O-Mad2:C-Mad2:Mad1^MIM^ dimer, we performed MD simulations on the full-length O-Mad2:C-Mad2:Mad1^MIM^ complex, modelled on the crystal structure of O-Mad2^LL^:C-Mad2:MBP1^MIM^ (Ref. ^16^). A total of 5.4 µs of simulations were carried out at 300 K. The Cα RMSFs of O-Mad2 in a O-Mad2:C-Mad2 dimer are highly comparable to that of monomeric O-Mad2 (Fig. 6e), except for small decreases at the C-terminus of the αC helix, located at the dimer interface (Fig. 6f). Notably, in contrast to O-Mad2, MD simulations indicated that the Cα RMSFs of both the β5-αC loop and residues 160-180 in C- Mad2 are substantially reduced, being equivalent to Cα residues of the core (Supplementary Fig. 10f). In C-Mad2, residues 160-180 constitute the ‘safety-belt’, whereas the β5-αC loop is restricted by the extended αA helix. The C-terminus of C-Mad2 is also substantially less dynamic than in O-Mad2, whereas the ten N-terminal residues remain highly dynamic in both states (Supplementary Fig. 10f). This agrees with our observation that the N-terminal residues do not adopt the αN helix in C-Mad2 in solution (Supplementary Fig. 3d). We conclude that despite the structural differences at the dimeric interface (Supplementary Fig. 10e), O-Mad2 retains similar structural and dynamic properties upon dimerisation and likely undergoes the same O- to C-Mad2 conversion pathway upon Cdc20^MIM^ binding.

### Cdc20^MIM^ promotes release of the β1 strand

The local differences in backbone dynamics (Fig. 4) suggested that different structural components in O-Mad2 might undergo conversion at different rates. To understand the molecular mechanism of the MIM-induced O- to C-Mad2 conversion, we recorded the conversion rates of individual residues by monitoring their change in peak intensity in time-resolved consecutive 2D NMR spectra (Fig. 7a). For this experiment, we compared the conversion rates of O-Mad2^R133A^ in the absence of MIM (Fig. 7b) and the O-Mad2^LL/R133A^ mutant in the presence of Cdc20^MIM-123-152^ (Fig. 7c). We used the O-Mad2^LL/R133A^ mutant for the Cdc20^MIM^-induced conversion time-course because of its reduced conversion rate: ∼250 min compared with <15 min for O-Mad2^R133A^ (Figs. 3b and 5), thus allowing sufficient NMR spectra to be recorded to define a time-course. The disappearance of the backbone resonances from O- Mad2 and the appearance of the backbone resonances for ‘empty’ C-Mad2 or C-Mad2:Cdc20^MIM^ were analysed by the changes in their peak intensities over 42 h. It was assumed that Mad2 converted from the open-to-closed state at the end of the measurement and there was no reverse conversion. The disappearance of backbone resonances from O-Mad2 reflected the transition from the initial open state, whereas the appearance of backbone resonances for C-Mad2 indicated the complete conversion to the closed state. A direct correlation between the disappearance rates of O-Mad2 resonances and appearance rates of C-Mad2 resonances would indicate the absence of intermediate states on conversion. No global unfolding and refolding was observed during Mad2 conversion, in agreement with proton-deuterium exchange experiments ^46^.

**Figure 7.**
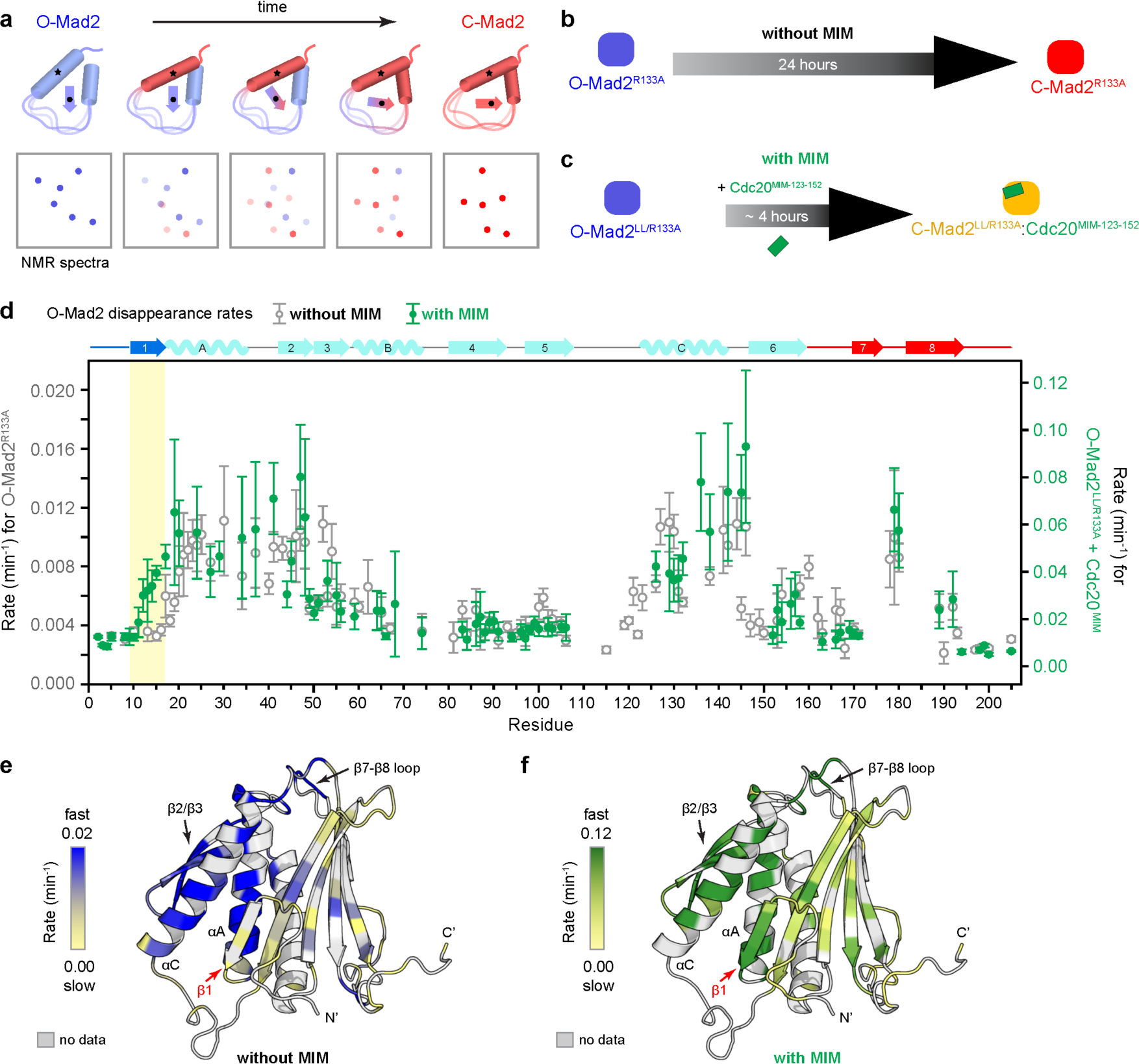
Cdc20^MIM^ promotes the release of the N-terminal β1 strand. a. Schematics illustrating the measurement of O- to C-Mad2 conversion rates using time-resolved NMR. A series of 2D HSQC spectra were collected at 25 °C for 42 h. The disappearance of backbone resonances from O-Mad2 reflects the transition from the initial open state, whereas the appearance of backbone resonances for ‘empty’ or ‘bound’ C-Mad2 indicates the complete conversion to the final closed state. **b, c.** Two samples were used for measurement: O-Mad2^R133A^ in the absence of Cdc20^MIM^ (**b**) and an O-Mad2^LL/R133A^ mutant that has a reduced conversion rate of ∼250 mins in the presence of Cdc20^MIM-123-152^ (**c**). **d.** The initial conversion rates of individual O-Mad2 residues were determined by plotting the peak intensities of their O-Mad2 resonances against time (Supplementary Fig. 11a, b) and fitting the data to an exponential decay curve. Error bars represent the standard regression errors of the fit. The rates for O-Mad2^R133A^ in the absence of Cdc20^MIM^ were plotted as grey open circles (without Cdc20^MIM^, left y-axis) and those for O-Mad2^LL/R133A^ in the presence of Cdc20^MIM-123-152^ were plotted as green circles (with Cdc20^MIM^, right y-axis). The data were scaled for comparison between the varying conversion rates across the sequence. The unscaled data are shown in Supplementary Fig. 11c. Distinct differences are observed for β1, highlighted in yellow. **e, f.** The initial conversion rates shown in (**d**) were mapped onto O-Mad2 in the absence of Cdc20^MIM^ (**e**) and in the presence of Cdc20^MIM^ ^123-152^ (**f**). The models are coloured in a scale, with blue (**e**) and green (**f)** showing the highest initial conversion rates. Residues with no available conversion data are coloured grey.

The disappearance of O-Mad2 resonances reflected the initial conversion rates with the data fitting closely to an exponential decay curve, both in the presence and absence of Cdc20^MIM^ (blue in Supplementary Fig. 11a, b). Different segments of O-Mad2^R133A^ showed distinct differences in their O-Mad2 disappearance rates (Fig. 7d). In particular, the αA helix, β2/β3 hairpin and αC helix exhibited the highest initial conversion rates. Despite the limited data available for the β7/β8 hairpin, signals from the β7-β8 loop indicated that the β7/β8 hairpin also underwent a high rate of conversion. O-Mad2^LL/R133A^ in the presence of Cdc20^MIM^ converted ∼8 times faster than O-Mad2^R133A^ without Cdc20^MIM^ (Supplementary Fig. 11c). However, both shared a mainly similar pattern of varying O-Mad2 disappearance rates across the sequence (Fig. 7d), suggesting O-Mad2 undergoes a very similar conversion pathway with or without MIM. However, a major difference was observed for the N-terminal β1 strand (Fig. 7d-f). The β1 strand was amongst the last to convert from its initial state in the absence of Cdc20^MIM^, but had a much higher initial conversion rate in the presence of Cdc20^MIM^. This indicated that Cdc20^MIM^ plays a role in triggering the conversion of the β1 strand, most likely by facilitating its release from the core.

While the resonances of O-Mad2 rapidly disappeared during Mad2 conversion, the resonances of C-Mad2 slowly appeared during conversion (red and orange in Supplementary Fig. 11a, b). The appearances of C-Mad2 resonances did not adopt an exponential pattern, unlike the disappearances of O-Mad2 resonances, and therefore could not be fitted to an exponential plateau curve. The lack of a direct correlation between the O-Mad2 disappearance rates and C- Mad2 appearance rates indicated there are one or more intermediates during the O- to C-Mad2 conversion. This agrees with our hypothesis that O-Mad2 has to undergo a partial conversion to fully convert to C-Mad2. This likely involves the unfolding of β7/β8 hairpin to expose its MIM- binding site, and the rearrangement of αC to facilitate β1 release.

## Discussion

SAC signalling, involving the kinetochore-catalysed conformational change of the metamorphic protein Mad2, represents an unusually complex regulatory mechanism. The conformational conversion of O-Mad2 to C-Mad2 is the rate limiting step in MCC assembly. Whereas the spontaneous Mad2 conversion rate *in vitro* is ∼24 h, Cdc20^MIM^ accelerates this conversion by >50-fold (Fig. 2c), effectively removing the kinetic energy barrier for MCC formation. Here, based on insights into the dynamics of structural elements of O-Mad2, and determination of conversion rates of individual residues on its transition to C-Mad2, we propose a model for how Cdc20^MIM^ induces Mad2 remodelling. Our work has implications for understanding the role of MCC-assembly catalysts, including those of the MCC-assembly scaffold, in facilitating this function of Cdc20^MIM^.

Our data confirm that the MIM-binding site is not accessible in O-Mad2 (Figs. 3 and 8a), in agreement with previous observations that Mad2 variants trapped in the open state do not bind Cdc20^MIM^ (Refs ^12,16,18,40^). For binding to occur, O-Mad2 must undergo a conformational conversion to expose the MIM-binding site that is obstructed by the β7/β8 hairpin. NMR dynamics data and MD simulations indicated that the β7/β8 hairpin is a highly flexible structural segment of O-Mad2 and may transiently unfold to expose the MIM-binding site (Figs. 4 and 8b). We showed that Cdc20^MIM^ binds with low affinity to a mutant of Mad2 lacking the β7/β8 hairpin (O-Mad2^Δβ7/β8^) (Fig. 3g), in agreement with a previous study ^40^. Since O-Mad2^Δβ7/β8^ is a structural mimic of O-Mad2 with a displaced β7/β8 hairpin, a likely crucial early event in the O- to C-Mad2 conversion is the displacement of the β7/β8 hairpin and pairing of Cdc20^MIM^ to β6, as proposed by Musacchio and colleagues ^28^. The low affinity of this interaction necessitates that Cdc20^MIM^ is positioned close, and in the correct relative orientation to its binding site on Mad2, a function fulfilled by the MCC-assembly scaffold that serves to increase the local concentrations of Cdc20^MIM^ and O-Mad2 ^31^. It also suggests that Cdc20^MIM^ is only locked in place in C-Mad2 by the ‘safety-belt’ created from a restructured β6-β7 loop, that is secured by the pairing of the β8’/β8” hairpin with β5 of the Mad2 core β sheet ^28^. The β8’/β8” hairpin in C-Mad2 is comprised of residues that form the β7/β8 hairpin and most of the flexible C-terminus of O- Mad2 (Fig. 1b, c). Pairing of the β8’/β8” hairpin with β5 requires displacement of β1 and its remodelling to form the N-terminus of αA. Without release of the β1 strand, the β7/β8 hairpin cannot be stably displaced and remodelled as β8’/β8”, and would compete with Cdc20^MIM^ for the MIM-binding site on β6 (Fig. 8c). Remodelling of β1 is slowed in the O-Mad2^LL^ mutant because the shortened β5-αC loop impedes the passage of β1-strand residues through the β5-αC loop, thereby restricting their access to the N-terminus of αA. In contrast, destabilising β1 as caused by the Mad2^L13A^ mutant, or its complete truncation as in Mad2^ΔN15^, traps Mad2 in a closed conformation ^16,43,44^. Thus, factors that promote β1 strand release and its remodelling would contribute to accelerating the O- to C-Mad2 conversion. In this study, we provide NMR evidence that Cdc20^MIM^ promotes the structural conversion of β1-strand residues from their O-Mad2 conformation (Fig. 7). This reveals a link between Cdc20^MIM^ binding and β1 strand displacement. One explanation for this is a potential allosteric coupling of β7/β8 hairpin and β1 strand release mediated through hydrophobic side chains on the β6 strand and αA and αC helices that link β7/β8 with β1 (Fig. 8f) ^16^. These conformational changes associated with the O-to C- Mad2 conversion might be triggered by loss of hydrophobic contacts between Phe186 of β8 and Phe151 of β6 caused by β7/β8 displacement. Phe151 in turn contacts the cluster of αA helix residues Phe23, Phe24 and Ile28. Rotation of the aromatic side chains of Phe23 and Phe24 blocks the binding pocket for Leu13 of β1 and is also linked to a shift of αC that contacts Ile11 of β1 (Fig. 8f). The rearrangement of aromatic residues is consistent with the distinctly different NMR fingerprint spectra for O-Mad2 and C-Mad2 (Supplementary Fig. 2d-e). Another factor linking Cdc20^MIM^ to β1 strand displacement is that Cdc20^MIM^ and the β7/β8 hairpin compete for binding to the β6 strand (Fig. 8c). With Cdc20^MIM^ bound to β6, residues of the displaced β7/β8 hairpin would be more likely to compete with β1 to pair with β5 (Fig. 8d, e). Transient displacement of the β7/β8 hairpin allows Cdc20^MIM^ binding, and lowers the free energy state of this MIM-bound intermediate (relative to non-bound). Stabilisation and formation of the newly folded C-Mad2 β8’/β8” hairpin and N-terminal turn of the αA helix is likely cooperative and coordinated. Hydrogen bonds linking Ser16 of αA with Thr188 and His191 of β8’/β8” stabilise C-Mad2 ^40,47^, whereas disruption of these hydrogen bonds was proposed to contribute to the ATP-dependent TRIP13-p31^comet^- catalysed remodelling of C-Mad2 into O-Mad2 ^48^.

**Figure 8.**
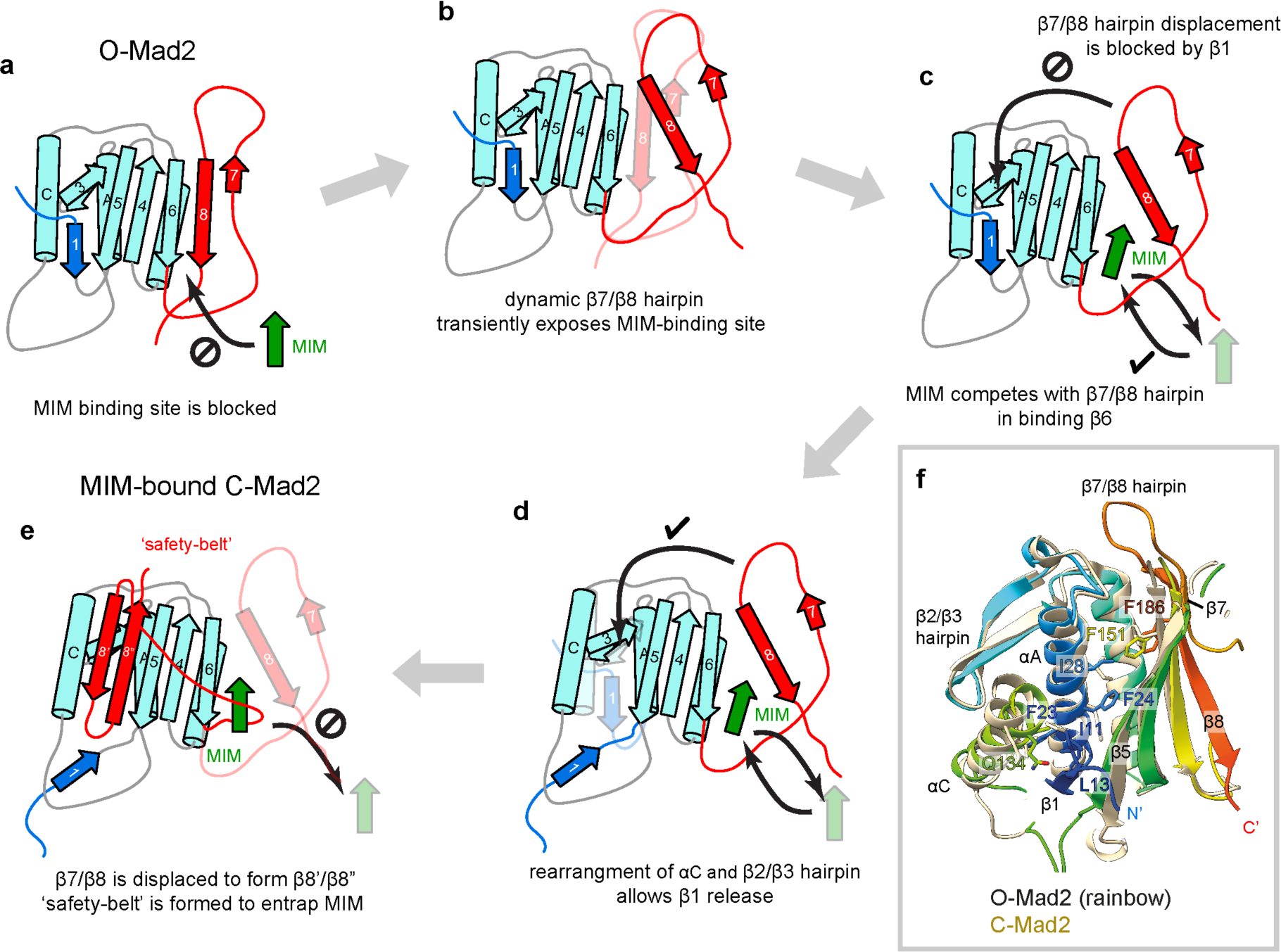
Molecular mechanism of Mad2 conversion. Secondary structure topology diagrams illustrating the molecular mechanism of the O- to C-Mad2 conversion. **a.** and **e.** show the topology of O- Mad2 and MIM-bound C-Mad2 respectively. Residues 1-10 of MIM-bound C-Mad2 is represented as unstructured. Displaced β-strands without hydrogen bonding to the Mad2-core are expected to be unstructured, but are shown here as β-strands for illustration. For simplicity, O-Mad2 is shown as a monomer, but a similar conversion mechanism is expected for O-Mad2 in an O:C dimer. **b.** The β7/β8 hairpin is highly dynamic and transiently unfolds to expose the MIM-binding site. **c.** Cdc20^MIM^ only weakly associates with β6 and competes with the β7/β8 hairpin for binding. **d.** Complete remodelling of β7/β8 is blocked by the β1 strand. **e.** Cdc20^MIM^ binding to β6 releases the β6/β7 hairpin allowing formation of the β8’/β8” hairpin that competes with β1 for pairing with β5. The reconfigured C-terminus of C-Mad2 entraps Cdc20^MIM^ through a ‘safety-belt’ connecting β6 with β8’. The transient Mad2:Cdc20^MIM^ intermediate has a lower free energy than the ‘empty’ Mad2 equivalent, reducing the energy barrier to C-Mad2 formation. **f.** Superimposition of O-Mad2 (N- to C-termini coloured blue-to- red) onto C-Mad2 (straw) (from O-Mad2:C-Mad2:MBP1^MIM^ (Ref. ^16^)) illustrating the network of hydrophobic residues linking Phe186 of β8 to Ile11 and Leu13 of β1 through Phe151 of β5, Ile28, Phe24 and Phe23 of αA. Side chains of these residues reposition, as does αC, on the O-to C-Mad2 conversion.

Finally, the self-association of O-Mad2 with C-Mad2 modestly contributes to catalysing the O- to C-Mad2 conversion, as originally proposed in the Mad2 template model ^18^ and demonstrated experimentally ^17–21,37^. As Mad2 dimerisation led to lower sensitivity and longer acquisition times in NMR measurements, we were unable to use NMR to demonstrate that dimerisation of O-Mad2 with C-Mad2 increases conversion rates induced by Cdc20^MIM^. Two previous studies reported differences in the HSQC spectra between monomeric O-Mad2 and O-Mad2 in the context of the O-Mad2:C-Mad2 dimer ^44,45^, indicative of dimer-induced structural changes mainly localised to interfacial residues of the αC helix and β2/β3 hairpin ^45^. O-Mad2 dimerisation with C-Mad2 may therefore induce structural changes at the β5/αC cleft of O-Mad2 contributing to destabilising β1, and explaining how dimerisation contributes to catalysing the O- to C-Mad2 conversion.

The capacity of Cdc20^MIM^ peptides to greatly accelerate the intrinsically slow O- to C-Mad2 conversion provides mechanistic insight into how unattached kinetochores activate the SAC ^1^. SAC signalling is mediated by the MCC ^2^, whose assembly is rate-limited by formation of C- Mad2:Cdc20. SAC catalysts ^20,27,28^ generate an MCC-assembly scaffold that functions to present the exposed Cdc20^MIM^ to its binding site on Mad2 ^27,28,30–32^. In contrast to the rapid association of Cdc20^MIM^ peptides to empty C-Mad2, full-length Cdc20 does not readily bind empty C-Mad2 ^28^, likely due to the steric hindrance of threading the long N-terminal segment of Cdc20 through the closed safety-belt of C-Mad2 ^28^. Thus, an inherent feature of SAC activation, rapid formation of C-Mad2, is coordinated to the presentation of its ligand, Cdc20^MIM^. This is achieved by the exposure of Cdc20^MIM^ through relief of a Cdc20-auto-inhibitory mechanism, combined with its close spatial positioning to its cryptic binding site on O-Mad2. Because empty C-Mad2 is non-productive for Cdc20 binding ^28^, the Cdc20^MIM^-induced triggering of the O- to C-Mad2 conversion is an elegant mechanism to regulate MCC assembly and thus SAC signalling.

Understanding how Cdc20^MIM^ directly induces the O- to C-Mad2 conversion is essential to our understanding of how MCC assembles. However, the intrinsic nature of O-Mad2 to undergo conversion to C-Mad2 has made it difficult to fully characterise this protein. Building on decades of structural work on Mad2, our study provides new insights into the dynamics of O-Mad2, demonstrating how protein dynamics coupled to Cdc20^MIM^ binding play a key mechanistic role in accelerating the Mad2 conversion rate, and serves as a framework for understanding the mechanisms of conformational conversion of metamorphic proteins.

## Methods

### Cloning Mad2 expression vectors

The cloning of human wild type Mad2 and the mutant Mad2^R133A^ by the USER® method (NEB) into the pRSFDuet-1 vector (71341-3, Sigma-Aldrich) with a His_6_-DoubleStrep-TEV tag at its N-terminus was described previously ^31^. For this study, Mad2 mutants were generated as follows.

The Mad2^LL^ expression construct was generated from the wild type Mad2 expression vector ^31^ by replacing residues 109-117 (TAKDDSAPR) with GSG. Mad2^ΔC10^ and Mad2^Δβ7/β8^ were modified from wild type Mad2 expression construct by deleting residues 196-205 and 166-205, respectively. Mad2^LL/R133A^ was modified by substituting Ala for Arg133 from the Mad2^LL^ expression construct. Mad2^LL/1′C10^ was modified from Mad2^LL^ by deleting residues 196-205.

Mad2^L13A/R133A^ was generated from the Mad2^R133A^ mutant. Site-directed mutagenesis of Mad2 was completed using the QuikChangeTM Lightning Site-Directed Mutagenesis Kit (Agilent) developed by Stratagene Inc. (La Jolla, CA). A schematic of Mad2 variants is shown in Fig. 1e.

### Protein expression and purification

Mad2 constructs were expressed in BL21 Star (DE3) cells (Thermo Fisher Scientific), co-transformed with a pRARE plasmid (Merck). Uniformly-labelled proteins were expressed in M9 minimal media (6 g/L Na_2_HPO_4_, 3 g/L KH_2_PO_4_, 0.5 g/L NaCl) supplemented with 1.7 g/L yeast nitrogen base without NH_4_Cl and amino acids (Sigma Y1251). 1 g/L ^15^NH_4_Cl and 4 g/L glucose were supplemented for ^15^N labelling. For ^13^C/^15^N double-labelled samples, unlabelled glucose was replaced with 3 g/L ^13^C-glucose. Protein expression was induced with 0.4 mM IPTG at 16 °C overnight. All samples were purified as previously described ^49^ and all O-Mad2 variants were kept at 4 °C during purification to prevent conversion to C-Mad2. Prior to all NMR experiments, proteins were dialysed into 20 mM HEPES, pH 7.0, 100 mM NaCl, 1 mM TCEP. ^13^C/^15^N-labelled samples were used for backbone assignment and all other experimental data were acquired on ^15^N-labelled samples. Proteins were kept on ice before NMR experiments to ensure all O-Mad2 remained in the open state. For ‘bound’ C-Mad2, MIM peptides were added in 1:2 molar excess to ensure complete conversion.

For wild-type Mad2, the protein was expressed as O-Mad2^WT^ using the conditions described above. C-Mad2^WT^ was generated by adding an excess of Mad1^MIM^ peptide to O-Mad2^WT^ and the sample was further purified using size-exclusion chromatography (SEC), where only monomeric C-Mad2^WT^:Mad1^MIM^ was used for analysis. The dimeric O-Mad2^WT^:C-Mad2^WT^:Mad1^MIM^ sample was prepared by mixing isotopically labelled O-Mad2^WT^ with a two-fold molar excess of unlabelled C-Mad2^WT^:Mad1^MIM^. The sample was further purified using SEC, where only dimeric O:Mad2^WT^:C-Mad2^WT^:Mad1^MIM^ was used for analysis.

### Peptide synthesis

All peptides were synthesised by Cambridge Research Biochemicals, UK. Peptides were dialysed into 20 mM HEPES, pH 7.0, 100 mM NaCl, 1 mM TCEP for NMR experiments. The sequences for the peptides used in this study are: Cdc20^MIM-111-138^ (EHQKAWALNLNGFDVEEAKILRLSGKPQ), Cdc20^MIM-123-137^ (FDVEEAKILRLSGKP), Cdc20^MIM-123-152^ (FDVEEAKILRLSGKPQNAPEGYQNRLKVLY), Cdc20^MIM-138-152^ (QNAPEGYQNRLKVLY), Mad1^MIM-529-550^ (RALQGDYDQSRTKVLHMSLNPT), MBP1^MIM^ (SWYSYPPPQRAV).

### Sample conditions for NMR

All NMR data on Mad2 were acquired in a 5 mm tube at 25 °C in 20 mM HEPES, pH 7.0, 100 mM NaCl, 1 mM TCEP, 5% D_2_O. Backbone assignment experiments and all spectra unless otherwise denoted were collected using an in-house Bruker 800 MHz Avance III spectrometer, equipped with a triple resonance TCI CryoProbe. 2D spectra were acquired using BEST ^1^H,^15^N- TROSY HSQC, with 16 scans and a recycle delay of 400 msec, giving an experimental time of 12 min per spectrum. A Bruker 950 MHz Avance III spectrometer (MRC Biomedical NMR Centre, Francis Crick Institute) was used for the residue conversion rate analysis and the characterisation of Mad2 dimers. Most experiments including all backbone experiments were acquired with 100 µM samples, the residue specific conversion rate analysis and characterisation of Mad2 dimers utilised 200 µM samples and 300 µM samples, respectively.

### Backbone assignment and secondary structure analysis

Backbone resonance assignments of multiple conformers of Mad2 were based on the following triple resonance experiments acquired with ^13^C,^15^N labelled samples: HNCO, HN(CA)CO, HNCA, HN(CO)CA, HNCACB, CBCA(CO)NH (Bruker pulse sequence library). All 3D datasets were collected with 15-30% non-uniform sampling and processed in MddNMR ^50^ using compressed sensing reconstruction. Backbone resonances were assigned in Sparky3 (T.D.Goddard & D.G.Kneller, UCSF) supported by in-house scripts and Mars ^51^. 2D ^1^H,^1^H NOESY spectra were recorded with mixing times of 100 ms to assign the indole NH resonances of the tryptophan side chains. Topspin 4.1.1 (Bruker) was used for processing and NMRFAM-Sparky 1.47 for data analysis ^52^.

Backbone assignment for full-length O-Mad2 was achieved using a combination of four variants, Mad2^R133A^ (a dimerisation-deficient mutant ^15^), Mad2^LL^ (Ref. ^16^), Mad2^ΔC10^ and Mad2^LL+ΔC10^.

Mad2^ΔC10^ and Mad2^LL+ΔC10^ provide the long-term stability required for NMR triple resonance experiments. The backbone resonances for the missing β5-αC loop in Mad2^LL^ were assigned using Mad2^ΔC10^. The backbone resonances for the C-terminal residues were assigned using a limited set of Cα chemical shifts from Mad2^LL^, as Mad2^LL^ also has a limited stability of <3 days and undergoes slow O- to-C Mad2 conversion. The HSQC spectra of these variants overlapped closely, indicating that the global folds of these mutants are similar (Supplementary Fig. 6a). All the assignments were consolidated and transferred to full-length Mad2^R133A^ (Supplementary Fig. 2a). Backbone resonances for ‘empty’ C-Mad2 were assigned using the double mutant Mad2^L13A/R133A^ (Ref. ^40^) (Supplementary Fig. 2b). Backbone resonances of C-Mad2:Cdc20^MIM^ were assigned using full-length Mad2^L13A/R133A^, in the presence of a two-fold molar excess of Cdc20^MIM^, spanning residues 111-138 (Cdc20^MIM-111-138^) (Supplementary Fig. 2c). Of the 199 non-proline residues in Mad2, the backbone amide resonances of 177 residues were assigned for O-Mad2, 163 residues were assigned for ‘empty’ C-Mad2 and 177 residues C-Mad2:Cdc20^MIM^. Tryptophan side chains resonances shown in Fig. 2a were assigned from ^1^H,^1^H NOESY experiments (O-Mad2, C-Mad2, C-Mad2:Cdc20^MIM^) or taken from the BMRB database (4775 for O-Mad2 and 5299 for C-Mad2:MBP1^MIM^). The combined data allowed the mapping of tryptophan side chain chemical shifts in the experimentally determined structures (PDB 1DUJ, 1S2H, 6TLJ, 1GO4 and 1KLQ) ^5,12-15,39^.

Secondary chemical shifts were calculated using the equation (δCɑ_obs_ – δCɑ_rc_) – (δCβ_obs_ – δCβ_rc_) where δCɑ_obs_ and δCβ_obs_ are the observed Cɑ and Cβ chemical shifts, and δCɑ_rc_ and δCβ_rc_ are the Cɑ and Cβ chemical shifts for random coils ^38,53^. Random coil chemical shifts were calculated based on sequence using POTENCI ^54^, taking into account the effects of pH, temperature and buffer pKa.

### Chemical shift perturbation (CSP) analysis

Weighted chemical shift perturbations were calculated using the equation ^55^: Δδ = [(Δδ_HN_W_HN_)^2^ + (Δδ_N_W_N_)^2^)^2^]^1/2^ where Δδ^1^H and Δδ^15^N are the chemical shift perturbations in ^1^H and ^15^N dimensions respectively ^56^. The weight factors were determined from the average variances of chemical shifts in the BMRB database ^57^, with W_HN_ = 1 and W_N_ = 0.16. Binding affinities K_D_ were estimated for residues with the most significant CSP. CSPs were plotted against the ligand concentration and fitted to the following equation: Δδ_obs_ = Δδ_max_ [(P+L+K_D_) – (P+L+K_D_)^2^ – 4PL]^1/2^ / 2P (Ref. ^55^) where P and L are the total concentrations of protein and ligand, respectively, Δδ_obs_ is the observed CSP and Δδ_max_ is the maximum CSP upon saturation.

### Relaxation measurements

_15_N T2 relaxation times of O-Mad2^ΔC10^ were measured using INEPT-based pseudo-3D pulse sequences with a recovery delay of 5 s incorporating a temperature-compensation scheme ( hsqct2etf3gpsitc3d, Bruker). Twelve mixing times were collected (8.48, 16.96, 33.92, 50.88, 67.84, 101.76, 135.68, 169.6, 203.52, 237.44, 271.36, and 8.48 ms). Data analysis was done in NMRFAM-Sparky 1.47 ^52^.

### SEC-MALS

Samples were analysed using a Heleos II 18-angle light scattering instrument (Wyatt Technology) and Optilab rEX online refractive index detector (Wyatt Technology) at 4 °C. 100 µl Mad2 sample at 300 µM was loaded onto a Superdex 75 10/300 GL increase column (GE Healthcare) pre-equilibrated with 20 mM HEPES, pH 7.0, 100 mM NaCl, 1 mM TCEP, 5 % D_2_O, running at 0.5 ml/min. The molecular mass was determined from the intercept of the Debye plot using the Zimm model as implemented in the Astra software (Wyatt Technology). Protein concentration was determined from the excess differential refractive index based on a 0.186 refractive index increment for 1 g/mL protein solution.

### Analysis of Mad2-residue conversion rates

Consecutive series of ^1^H,^15^N SOFAST-HMQC experiments were acquired in a pseudo-3D fashion to ensure all spectra were collected in near-identical conditions. SOFAST-HMQC was chosen for its short pulse sequence to maximise the number of data points collected during Mad2 conversion. All parameters were set up using a test sample to minimise the dead-time in data acquisition. Each 2D spectrum was acquired with eight scans and a recycle delay of 200 msec and 256 complex points in the indirect dimension, giving a final spectral resolution of 2.8 Hz per points in the indirect dimension and an experimental time of 5 min per spectrum. A total of 500 spectra were collected to monitor Mad2 conversion over 42 h. All spectra were processed using in-house scripts and NMRPipe ^58^. Four spectra were sequentially added to enhance signal-to-noise, and consecutive data points have a 5-min offset. Peak intensities for each added spectrum were analysed using Poky build:20230213 ^59^ and extracted using a modified Python module.

Peak intensities for O-Mad2 were normalised with the initial peak intensities in the first added spectrum, assuming all Mad2 molecules were in the open state at the initial time point. Peak intensities for C-Mad2 were normalised with the final peak intensities in the last added spectrum. It was assumed that all Mad2 proteins were converted from open to closed at the end of the measurement and there was no reverse conversion. For the disappearance of O-Mad2 resonances, data were fitted to an exponential decay curve with nonlinear regression, using the equation: y = (y_0_ – y_min_) exp (-*k*x) + y_min_, where y_0_ is the initial peak intensity, y_m_ is the minimum peak intensity and *k* is the rate constant. For the appearance of C-Mad2 resonances, the data were not fitted as they do not show a clear exponential or delayed exponential increase. Peaks that are overlapping in the O-Mad2 and C-Mad2 spectra were discarded in the analysis. Normalisation of peak intensities, curve fitting and plotting were carried out using in-house python scripts.

### MD simulations

The starting structure for MD simulations on monomeric full-length O-Mad2 was built using the crystal structure of O-Mad2^LL^ (PDB 2V64 ^16^, chain D) and the missing segments (residues 1-8, 109-118, 195-205) were built using Modeller ^60^. The C-terminal 10 residues were manually truncated for the simulation of Mad2^ΔC10^. The starting structure for MD simulations on full-length O-Mad2:C-Mad2:Mad1^MIM^ was built using the crystal structure of O-Mad2^LL^:C- Mad2:MBP1^MIM^ (PDB 2V64 ^16^). The missing segments in O-Mad2 were built as previously described and the MBP1^MIM^ was manually modified to match the sequence of Mad1^MIM-538-551^ (SRTKVLHMSLNPTS) in PyMOL (Schrödinger, LLC), guided by the structure of C- Mad2^R133A^:Mad1^485-718^ (PDB 1GO4 ^15^). Cys79 and Cys106 were defined as unoxidised. 100 mM NaCl was added to the solvent. Simulations were carried out using the DES-Amber force field and TIP4P-D water model ^61^. The DES-Amber force field was chosen for its suitability to study proteins with both folded and disordered regions. MD simulations were carried out in GROMACS ^62^ with the Verlet leapfrog integrator, a 2 fs time step, constrained bonds to hydrogen, Particle Mesh Ewald for long-range electrostatics and a 1 nm cut-off for non-bonded interactions. A 100 ps NVT equilibration and a 100 ps NPT equilibration preceded 0.9 μs production runs in the NPT ensemble, with snapshots saved every 200 ps. A total of 5.4 µs of simulations were carried out for each Mad2 variant or at each temperature, using six independent simulations of 0.9 µs. For each snapshot the RMSD was calculated for all protein backbone atoms to the starting structure. Cα RMSF was calculated by fitting each snapshot to the average coordinates, calculating the mean squared distance to the average coordinates across snapshots for each Cα, and averaging over the simulation repeats. Both RMSD and RMSF were calculated using standard GROMACS ^62^ scripts and plotted using GraphPad PRISM 9.5.1.

### Other computational methods

Molecular graphics were produced in PyMOL Molecular Graphics System, Version 2.5.3 Schrödinger, LLC. Secondary structure topology diagrams were drawn with Biotite ^63^ and CCP4 Topdraw ^64^, using their structural annotations on the NCBI database.

### Data Availability

The backbone assignments in this study were deposited to the BMRB database (http://www.bmrb.wisc.edu/), with the accession numbers: 52275 for O-Mad2^FL^, 52276 for ‘empty’ C-Mad2^L13A/R133A^ and 52277 for C-Mad2^R133A^:Cdc20^MIM^. Simulation trajectories are available at Zenodo (https://doi.org/10.5281/zenodo.10557066).

## Acknowledgements

This work was funded by MRC grant (MC_UP_1201/6) and CRUK grant (C576/A14109) to D.B. and MRC grant (MC_UP_1201/33) to J.G.G. E.F. was funded by a Gates Cambridge Scholarship. We thank Chris Johnson for useful discussion on protein folding and Chris Batters for his technical support on SEC-MALS. NMR studies at 950 MHz were supported by the Francis Crick Institute through access to the MRC Biomedical NMR Centre. The Francis Crick Institute receives its core funding from Cancer Research UK (FC001029), the UK Medical Research Council (FC001029), and the Wellcome Trust (FC001029). We thank Geoff Kelly (Francis Crick Institute) for his advice and technical support. For the purpose of open access, the authors have applied a CC BY public copyright license to any Author Accepted Manuscript version arising.

## Author Contributions

C.Y., E.F. and D.B. designed the experiments. E.F., C.Y., J.Y. and Z.Z. prepared the samples. C.Y. and S.F. acquired and analysed the NMR data. J.G.G and C.Y. set up and analysed the MD simulations. C.Y. performed the SEC-MALS. C.Y. and D.B. wrote the manuscript with input from all authors.

## References

1 Rieder, C. L., Cole, R. W., Khodjakov, A. & Sluder, G. The checkpoint delaying anaphase in response to chromosome monoorientation is mediated by an inhibitory signal produced by unattached kinetochores. The Journal of cell biology 130, 941–948 (1995).

2 Sudakin, V., Chan, G. K. & Yen, T. J. Checkpoint inhibition of the APC/C in HeLa cells is mediated by a complex of BUBR1, BUB3, CDC20, and MAD2. The Journal of cell biology 154, 925-936 (2001).

3 Chao, W. C., Kulkarni, K., Zhang, Z., Kong, E. H. & Barford, D. Structure of the mitotic checkpoint complex. Nature 484, 208–213, doi:nature10896 [pii] 10.1038/nature10896 (2012).

4 Izawa, D. & Pines, J. The mitotic checkpoint complex binds a second CDC20 to inhibit active APC/C. Nature 517, 631–634, doi:10.1038/nature13911 (2015).

5 Alfieri, C., et al. Molecular basis of APC/C regulation by the spindle assembly checkpoint. Nature 536, 431–436, doi:10.1038/nature19083 (2016).

6 Yamaguchi, M. et al. Cryo-EM of Mitotic Checkpoint Complex-Bound APC/C Reveals Reciprocal and Conformational Regulation of Ubiquitin Ligation. Molecular cell 63, 593–607, doi:10.1016/j.moticel.2016.07.003 (2016).

7 Musacchio, A. The Molecular Biology of Spindle Assembly Checkpoint Signaling Dynamics. Current biology : CB 25, R1002–1018, doi:10.1016/j.cub.2015.08.051 (2015).

8 Fischer, E. S. Kinetochore-catalyzed MCC formation: A structural perspective. IUBMB Life 75, 289–310, doi:10.1002/iub.2697 (2023).

9 McAinsh, A. D. & Kops, G. Principles and dynamics of spindle assembly checkpoint signalling. Nature reviews. Molecular cell biology 24, 543–559, doi:10.1038/s41580-023-00593-z (2023).

10 Watson, E. R., Brown, N. G., Peters, J. M., Stark, H. & Schulman, B. A. Posing the APC/C E3 Ubiquitin Ligase to Orchestrate Cell Division. Trends in cell biology 29, 117–134, doi:10.1016/j.tcb.2018.09.007 (2019).

11 Barford, D. Structural interconversions of the anaphase-promoting complex/cyclosome (APC/C) regulate cell cycle transitions. Current opinion in structural biology 61, 86–97, doi:10.1016/j.sbi.2019.11.010 (2020).

12 Luo, X. et al. Structure of the Mad2 spindle assembly checkpoint protein and its interaction with Cdc20. Nat Struct Biol 7, 224–229, doi:10.1038/73338 (2000).

13 Luo, X., Tang, Z., Rizo, J. & Yu, H. The Mad2 spindle checkpoint protein undergoes similar major conformational changes upon binding to either Mad1 or Cdc20. Molecular cell 9, 59–71 (2002).

14 Luo, X. et al. The Mad2 spindle checkpoint protein has two distinct natively folded states. Nature structural & molecular biology 11, 338–345, doi:10.1038/nsmb748 (2004).

15 Sironi, L. et al. Crystal structure of the tetrameric Mad1-Mad2 core complex: implications of a ’safety belt’ binding mechanism for the spindle checkpoint. The EMBO journal 21, 2496–2506 (2002).

16 Mapelli, M., Massimiliano, L., Santaguida, S. & Musacchio, A. The Mad2 conformational dimer: structure and implications for the spindle assembly checkpoint. Cell 131, 730–743 (2007).

17 Lad, L., Lichtsteiner, S., Hartman, J. J., Wood, K. W. & Sakowicz, R. Kinetic analysis of Mad2-Cdc20 formation: conformational changes in Mad2 are catalyzed by a C-Mad2- ligand complex. Biochemistry 48, 9503–9515, doi:10.1021/bi900718e (2009).

18 De Antoni, A. et al. The Mad1/Mad2 complex as a template for Mad2 activation in the spindle assembly checkpoint. Current biology : CB 15, 214–225, doi:S0960982205000734 [pii] 10.1016/j.cub.2005.01.038 (2005).

19 SimoneNa, M. et al. The influence of catalysis on mad2 activation dynamics. PLoS biology 7, e10, doi:10.1371/journal.pbio.1000010 (2009).

20 Faesen, A. C. et al. Basis of catalytic assembly of the mitotic checkpoint complex. Nature 542, 498–502, doi:10.1038/nature21384 (2017).

21 Kulukian, A., Han, J. S. & Cleveland, D. W. UnaNached kinetochores catalyze production of an anaphase inhibitor that requires a Mad2 template to prime Cdc20 for BubR1 binding. Developmental cell 16, 105–117 (2009).

22 Aravind, L. & Koonin, E. V. The HORMA domain: a common structural denominator in mitotic checkpoints, chromosome synapsis and DNA repair. Trends Biochem Sci 23, 284–286, doi:10.1016/s0968-0004(98)01257-2 (1998).

23 Gu, Y., Desai, A. & CorbeN, K. D. Evolutionary Dynamics and Molecular Mechanisms of HORMA Domain Protein Signaling. Annual review of biochemistry 91, 541–569, doi:10.1146/annurev-biochem-090920-103246 (2022).

24 Meraldi, P., Draviam, V. M. & Sorger, P. K. Timing and checkpoints in the regulation of mitotic progression. Developmental cell 7, 45–60, doi:10.1016/j.devcel.2004.06.006 (2004).

25 Clute, P. & Pines, J. Temporal and spatial control of cyclin B1 destruction in metaphase. Nature cell biology 1, 82–87 (1999).

26 Dick, A. E. & Gerlich, D. W. Kinetic framework of spindle assembly checkpoint signalling. Nature cell biology 15, 1370–1377, doi:10.1038/ncb2842 (2013).

27 Ji, Z., Gao, H., Jia, L., Li, B. & Yu, H. A sequential multi-target Mps1 phosphorylation cascade promotes spindle checkpoint signaling. eLife 6, doi:10.7554/eLife.22513 (2017).

28 Piano, V. et al. CDC20 assists its catalytic incorporation in the mitotic checkpoint complex. Science 371, 67–71, doi:10.1126/science.abc1152 (2021).

29 Fischer, E. S. et al. Molecular mechanism of Mad1 kinetochore targeting by phosphorylated Bub1. EMBO reports 22, e52242, doi:10.15252/embr.202052242 (2021).

30 Lara-Gonzalez, P., Kim, T., Oegema, K., CorbeN, K. & Desai, A. A tripartite mechanism catalyzes Mad2-Cdc20 assembly at unaNached kinetochores. Science 371, 64-+, doi:10.1126/science.abc1424 (2021).

31 Fischer, E. S. et al. Juxtaposition of Bub1 and Cdc20 on phosphorylated Mad1 during catalytic mitotic checkpoint complex assembly. Nature communicaJons 13, 6381, doi:10.1038/s41467-022-34058-2 (2022).

32 Chen, C. et al. The structural flexibility of MAD1 facilitates the assembly of the Mitotic Checkpoint Complex. Nature communicaJons 14, 1529, doi:10.1038/s41467-023-37235-z (2023).

33 Hiruma, Y. et al. CELL DIVISION CYCLE. Competition between MPS1 and microtubules at kinetochores regulates spindle checkpoint signaling. Science 348, 1264–1267, doi:10.1126/science.aaa4055 (2015).

34 Ji, Z., Gao, H. & Yu, H. CELL DIVISION CYCLE. Kinetochore aNachment sensed by competitive Mps1 and microtubule binding to Ndc80C. Science 348, 1260–1264, doi:10.1126/science.aaa4029 (2015).

35 Han, J. S. et al. Catalytic assembly of the mitotic checkpoint inhibitor BubR1-Cdc20 by a Mad2-induced functional switch in Cdc20. Molecular cell 51, 92–104, doi:10.1016/j.moticel.2013.05.019 (2013).

36 Zhang, Y. & Lees, E. Identification of an overlapping binding domain on Cdc20 for Mad2 and anaphase-promoting complex: model for spindle checkpoint regulation. Mol Cell Biol 21, 5190–5199, doi:10.1128/MCB.21.15.5190-5199.2001 (2001).

37 Nezi, L. et al. Accumulation of Mad2-Cdc20 complex during spindle checkpoint activation requires binding of open and closed conformers of Mad2 in Saccharomyces cerevisiae. The Journal of cell biology 174, 39–51, doi:10.1083/jcb.200602109 (2006).

38 Spera, S. & Bax, A. Empirical Correlation between Protein Backbone Conformation and C- Alpha and C-Beta C-13 Nuclear-Magnetic-Resonance Chemical-Shios. Journal of the American Chemical Society 113, 5490–5492, doi:DOI 10.1021/ja00014a071 (1991).

39 Alfieri, C., Tischer, T. & Barford, D. A unique binding mode of Nek2A to the APC/C allows its ubiquitination during prometaphase. EMBO reports 21, e49831, doi:10.15252/embr.201949831 (2020).

40 Yang, M. et al. Insights into mad2 regulation in the spindle checkpoint revealed by the crystal structure of the symmetric mad2 dimer. PLoS biology 6, e50 (2008).

41 Fang, G., Yu, H. & Kirschner, M. W. The checkpoint protein MAD2 and the mitotic regulator CDC20 form a ternary complex with the anaphase-promoting complex to control anaphase initiation. Genes & development 12, 1871–1883 (1998).

42 Zhao, Y. Y. et al. Stable folding intermediates prevent fast interconversion between the closed and open states of Mad2 through its denatured state. Protein Eng Des Sel 29, 23–29, doi:10.1093/protein/gzv056 (2016).

43 Yang, M. et al. p31comet blocks Mad2 activation through structural mimicry. Cell 131, 744–755, doi:S0092-8674(07)01199-3 [pii] 10.1016/j.cell.2007.08.048 (2007).

44 Mapelli, M. et al. Determinants of conformational dimerization of Mad2 and its inhibition by p31comet. The EMBO journal 25, 1273–1284, doi:7601033 [pii]10.1038/sj.emboj.7601033 (2006).

45 Hara, M., Ozkan, E., Sun, H., Yu, H. & Luo, X. Structure of an intermediate conformer of the spindle checkpoint protein Mad2. Proceedings of the NaJonal Academy of Sciences of the United States of America 112, 11252–11257, doi:10.1073/pnas.1512197112 (2015).

46 Brutione, M. L. et al. Mechanistic insight into TRIP13-catalyzed Mad2 structural transition and spindle checkpoint silencing. Nature communicaJons 8, 1956, doi:10.1038/s41467-017-02012-2 (2017).

47 Ye, Q. et al. The AAA+ ATPase TRIP13 remodels HORMA domains through N-terminal engagement and unfolding. The EMBO journal 36, 2419–2434, doi:10.15252/embj.201797291 (2017).

48 Alfieri, C., Chang, L. & Barford, D. Mechanism for remodelling of the cell cycle checkpoint protein MAD2 by the ATPase TRIP13. Nature 559, 274–278, doi:10.1038/s41586-018-0281-1 (2018).

49 Luo, X. & Yu, H. Purification and assay of Mad2: a two-state inhibitor of anaphase- promoting complex/cyclosome. Methods in enzymology 398, 246–255, doi:10.1016/S0076-6879(05)98020-8 (2005).

50 Jaravine, V. A., Zhuravleva, A. V., Permi, P., Ibraghimov, I. & Orekhov, V. Y. Hyperdimensional NMR spectroscopy with nonlinear sampling. J Am Chem Soc 130, 3927–3936, doi:10.1021/ja077282o (2008).

51 Jung, Y. S. & ZwecksteNer, M. Mars -- robust automatic backbone assignment of proteins. J Biomol NMR 30, 11–23, doi:10.1023/B:JNMR.0000042954.99056.ad (2004).

52 Lee, W., Tonelli, M. & Markley, J. L. NMRFAM-SPARKY: enhanced sooware for biomolecular NMR spectroscopy. BioinformaJcs 31, 1325–1327, doi:10.1093/bioinformatics/btu830 (2015).

53 Kakeshpour, T. et al. A lowly populated, transient beta-sheet structure in monomeric Abeta(1-42) identified by multinuclear NMR of chemical denaturation. Biophys Chem 270, 106531, doi:10.1016/j.bpc.2020.106531 (2021).

54 Nielsen, J. T. & Mulder, F. A. A. POTENCI: prediction of temperature, neighbor and pH- corrected chemical shios for intrinsically disordered proteins. J Biomol NMR 70, 141–165, doi:10.1007/s10858-018-0166-5 (2018).

55 Williamson, M. P. Using chemical shio perturbation to characterise ligand binding. Prog Nucl Magn Reson Spectrosc 73, 1–16, doi:10.1016/j.pnmrs.2013.02.001 (2013).

56 Ayed, A. et al. Latent and active p53 are identical in conformation. Nat Struct Biol 8, 756–760, doi:10.1038/nsb0901-756 (2001).

57 Mulder, F. A., Schipper, D., BoN, R. & Boelens, R. Altered flexibility in the substrate- binding site of related native and engineered high-alkaline Bacillus subtilisins. Journal of molecular biology 292, 111–123, doi:10.1006/jmbi.1999.3034 (1999).

58 Delaglio, F. et al. NMRPipe: a multidimensional spectral processing system based on UNIX pipes. J Biomol NMR 6, 277–293 (1995).

59 Lee, W., Rahimi, M., Lee, Y. & Chiu, A. POKY: a sooware suite for multidimensional NMR and 3D structure calculation of biomolecules. BioinformaJcs 37, 3041–3042, doi:10.1093/bioinformatics/btab180 (2021).

60 Webb, B. & Sali, A. Comparative Protein Structure Modeling Using MODELLER. Curr Protoc Protein Sci 86, 2 9 1–2 9 37, doi:10.1002/cpps.20 (2016).

61 Piana, S., Robustelli, P., Tan, D., Chen, S. & Shaw, D. E. Development of a Force Field for the Simulation of Single-Chain Proteins and Protein-Protein Complexes. J Chem Theory Comput 16, 2494–2507, doi:10.1021/acs.jctc.9b00251 (2020).

62 Abraham, M. et al. GROMACS: High performance molecular simulations through mulL- level parallelism from laptops to supercomputers. SoNwareX 1-2, 19-25, doi:10.1016/j.soox.2015.06.001 (2015).

63 Kunzmann, P. & Hamacher, K. Biotite: a unifying open source computational biology framework in Python. BMC bioinformaJcs 19, 346, doi:10.1186/s12859-018-2367-z (2018).

64 Bond, C. S. TopDraw: a sketchpad for protein structure topology cartoons. BioinformaJcs 19, 311–312, doi:10.1093/bioinformatics/19.2.311 (2003).

