## Supplementary Figures 1 to 11 for "Molecular mechanism of Mad2 conformational conversion promoted by the Mad2-interaction motif of Cdc20"

O-Mad2

| Construct | PDB | Missing residues | State | References |
|---|---|---|---|---|
| O-Mad2$^{LL}$ | 2V64 | 1-8, 109-118, 195-205 | O-C dimer | Mapelli *et al.* (2007) Ref. 16 |
| * O-Mad2$^{\Delta N10}$ | 1DUJ | 1-10, **a** 196-205 | monomer | Luo *et al.* (2000) Ref. 12 |
| **b** O-Mad2$^{\Delta N10}$ | 3GMH | 1-10, 204-205 | O-C dimer | Hara *et al.* (2015) Ref. 45 |

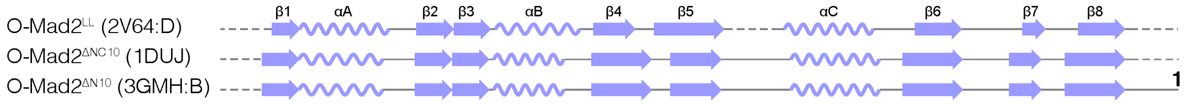


C-Mad2

| Construct | PDB | Missing residues | State | References |
|---|---|---|---|---|
| * 'empty' C-Mad2$^{R133A}$ | 1S2H | Full-length | monomer | Luo *et al.* (2004) Ref. 14 |
| C-Mad2$^{L13A/C79S/C106S}$ | 2VFX | Full-length | C-C dimer | Yang *et al.* (2008) Ref. 40 |

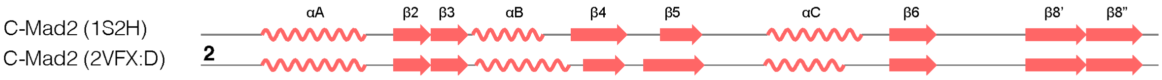


MIM-bound C-Mad2

| Construct | PDB | Missing residues | State | References |
|---|---|---|---|---|
| C-Mad2 + Cdc20 | 6TLJ | 1-7, 203-205 | monomer | Alfieri *et al.* (2016, 2020) Ref.5, 39 |
| C-Mad2 + MBP1 | 2V64 | Full-length | O-C dimer | Mapelli *et al.* (2007) Ref. 16 |
| * C-Mad2$^{\Delta N10}$ + MBP1 | 1KLQ | 1-10 | monomer | Luo *et al.* (2002) Ref. 13 |
| C-Mad2$^{R133A}$ + Mad1 | 1GO4 | 1-7, 204-205 | monomer | Sironi *et al.* (2002) Ref. 15 |
| C-Mad2$^{L13A}$ + MBP1 | 2QYF | 1-8 | dimer **c** | Yang *et al.* (2007) Ref. 43 |

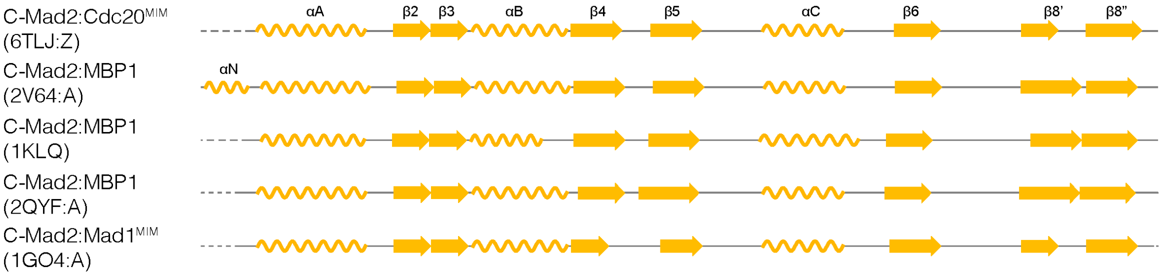


* Solution structures

**a** Residues 196-205 were excluded in structure determination
**b** O-Mad2$^{\Delta N10}$ dimerised to 'empty' C-Mad2$^{\Delta N10}$, named I-Mad2 in the paper
**c** MBP1-bound C-Mad2 forms a dimer with p31$^{comet}$, which mimics O-Mad2:C-Mad2 dimer

Features due to crystal packing:
**1** Residue 196-203 was inserted into the MIM binding site of another Mad2 O-C dimer
**2** Residue 1-9 was inserted into the MIM binding site of another Mad2 C-C dimer

**Supplementary Figure 1. Summary of structural studies on human Mad2.** A summary of structural work on human Mad2, highlighting the Mad2 variants used in each study. The oligomeric state listed refers to the interaction between Mad2 molecules, e.g. a Mad1:Mad2 complex is referred as monomeric Mad2 provided that the Mad2 in the structure does not form a dimer with another molecule of Mad2. A secondary structure topology diagram for each structure was generated based on their annotations in the NCBI database. O-Mad2 is represented in blue, 'empty' C-Mad2 in red, and MIM-bound C-Mad2 in orange. Solution structures are denoted with an asterisk (*). The crystal structure of O-Mad2$^{\Delta N10}$ was determined as a dimer bound to 'empty' C-Mad2$^{\Delta N10}$ (PDB 3GMH) [45] and referred to as intermediate Mad2 (I-Mad2) [45]. We refrain from designating any Mad2 conformation as an intermediate. A disulphide bond between residue 79 and 106 was observed in some crystal and cryo-EM structures.

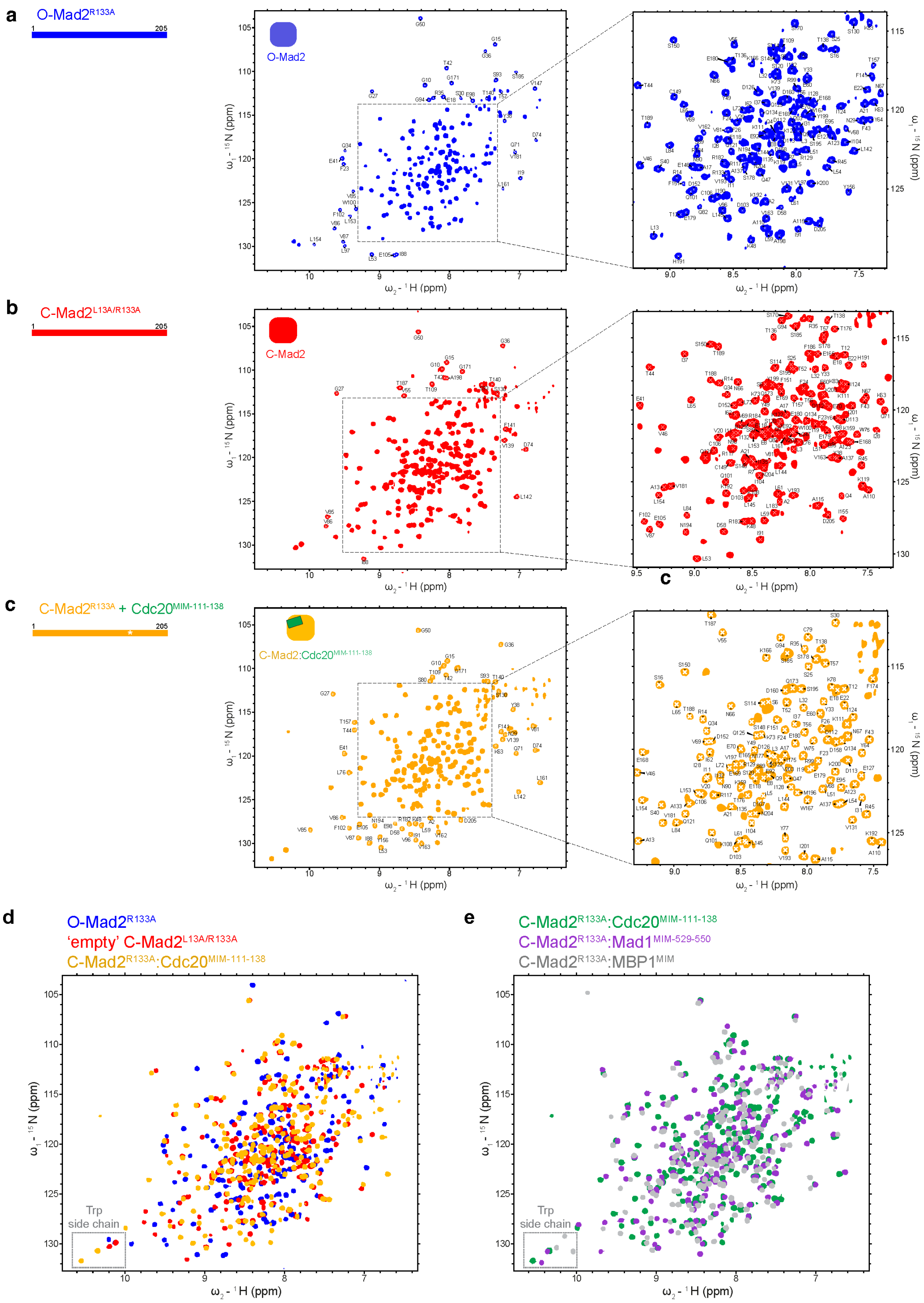


**Supplementary Figure 2. Backbone resonances of different Mad2 conformers.** HSQC spectra of **a.** full-length Mad2$^{R133A}$, **b.** 'empty' C-Mad2$^{L13A/R133A}$ and **c.** Cdc20$^{MIM-111-138}$-bound C-Mad2$^{R133A}$ are shown with the assignment of their backbone resonances. Of the 199 non-proline residues in Mad2, the backbone amide resonances of 177 residues were assigned for O-Mad2, 163 residues were assigned for 'empty' C-

Mad2 and 177 residues C-Mad2:Cdc20$^{MIM}$. **a.** Backbone assignments for full-length O-Mad2 were achieved using a combination of four Mad2 variants: Mad2$^{R133A}$, Mad2$^{LL}$, Mad2$^{\Delta C10}$ and Mad2$^{LL/\Delta C10}$. All assignments were consolidated and transferred to full-length Mad2$^{R133A}$. **b.** Backbone resonances for 'empty' C-Mad2 were assigned using the double mutant Mad2$^{L13A/R133A}$. **c.** Backbone resonances of C-Mad2:Cdc20$^{MIM}$ were assigned using full-length Mad2$^{R133A}$, in the presence of a two-fold molar excess of Cdc20$^{MIM\text{-}111\text{-}138}$ peptide to ensure complete open-to-closed conversion. NMR spectra were previously shown in [31] without backbone assignment. **d.** HSQC spectrum of 100 μM full-length O-Mad2$^{R133A}$ (blue), C-Mad2$^{L13A/R133A}$ (red) and C-Mad2$^{R133A}$:Cdc20$^{MIM\text{-}111\text{-}138}$ (orange). Reproduced from Fig. 7 of [31]. **e.** HSQC of C-Mad2$^{R133A}$ in the presence of a two-fold molar excess (200 μM) Cdc20$^{MIM\text{-}111\text{-}138)}$ (green), Mad1$^{MIM\text{-}529\text{-}550}$ (purple) and Mad1-binding peptide (MBP1$^{MIM}$) (grey). Tryptophan side-chain resonances were used to monitor the conformational changes in Mad2 and are boxed on the spectra.

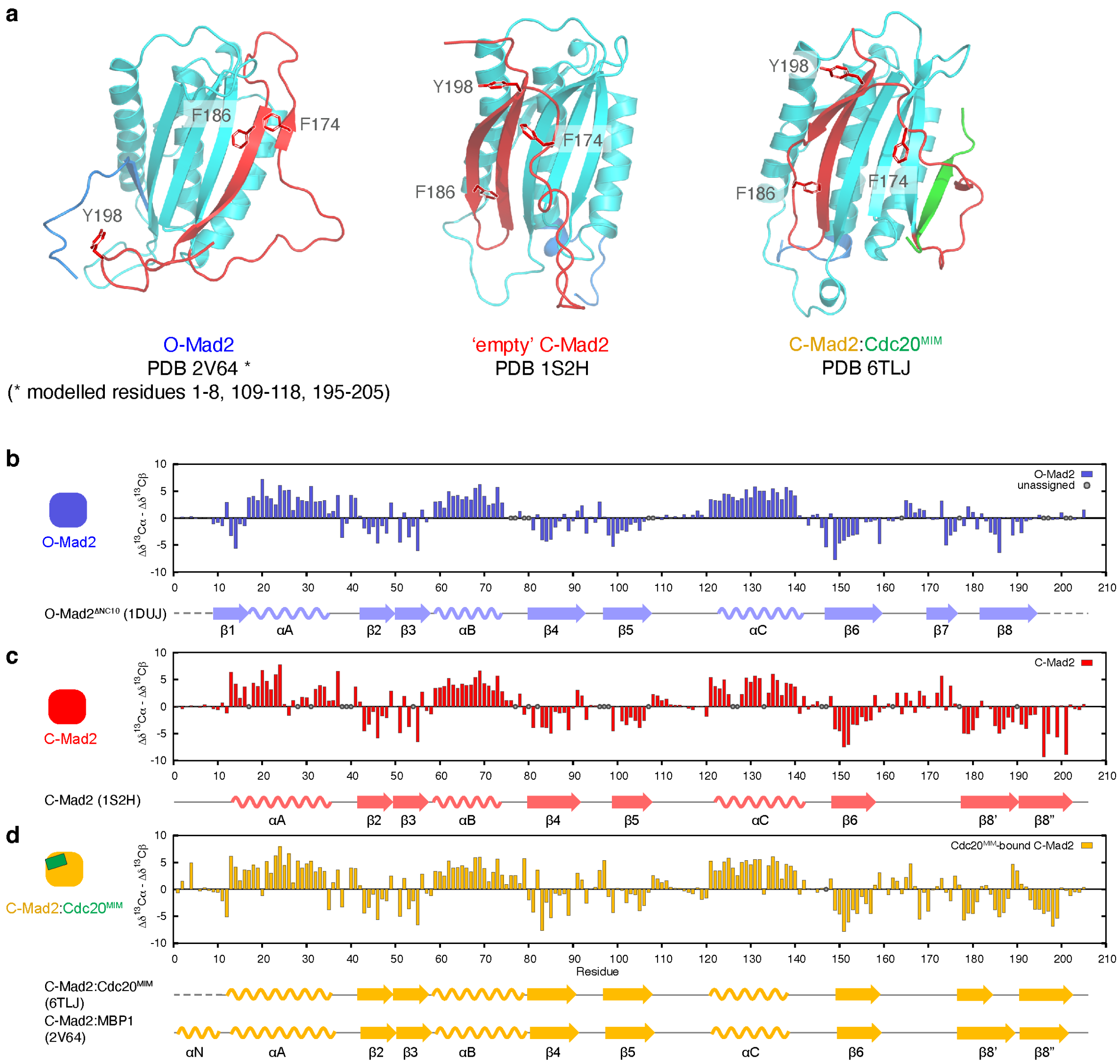


**Supplementary Figure 3. Solution secondary structure of different Mad2 conformers.** **a.** Representative structural models of Mad2 in different conformations. The O-Mad2 structure is based on the crystal structure of O-Mad2$^{LL}$ (PDB 2V64, chain D) [16] and the missing segments (residues 1-8, 109-118, 195-205) were built using Modeller [60]. The C-Mad2 structure is based on a solution structure of 'empty' C-Mad2 (PDB 1S2H) [14]. The structure of C-Mad2:Cdc20$^{MIM}$ is extracted from the cryo-EM structure of the APC/C:MCC (PDB 6TLJ) [5,39]. In O-Mad2, Phe174 of β7 and Phe186 of β8 are both buried within the hydrophobic core, whereas Tyr198, located near the C-terminus, is solvent exposed. Upon conversion, Phe174 becomes exposed as it forms part of the 'safety-belt', whereas Phe186 and Tyr198 become buried in their new positions in β8'/β8" as part of an extended antiparallel β sheet. Such a rearrangement manifests in substantial chemical shift changes for these aromatic amino acids and other nearby residues. **b-d**. Secondary chemical shifts ($\Delta\delta^{13}C\alpha - \Delta\delta^{13}C\beta$) reveal the secondary structure of Mad2 in solution. Positive values suggest α-helical propensity and negative values suggest propensity for β-strand conformation. Unassigned resonances are denoted as grey circles. Secondary structure topology diagrams from corresponding Mad2 structures are included for reference and residues absent in the structures are represented by dotted lines. Secondary chemical shifts for full-length O-Mad2$^{R133A}$, 'empty' C-Mad2$^{L13A/R133A}$ and C-Mad2$^{R133A}$:Cdc20$^{MIM-111-138}$ are shown in **b, c** and **d** respectively. **d.** Secondary chemical shifts of C-Mad2$^{R133A}$: Cdc20$^{MIM-111-138}$ indicates the absence of αN in solution. αN was observed in MBP1$^{MIM}$-bound C-Mad2 (PDB 2V64) [16].

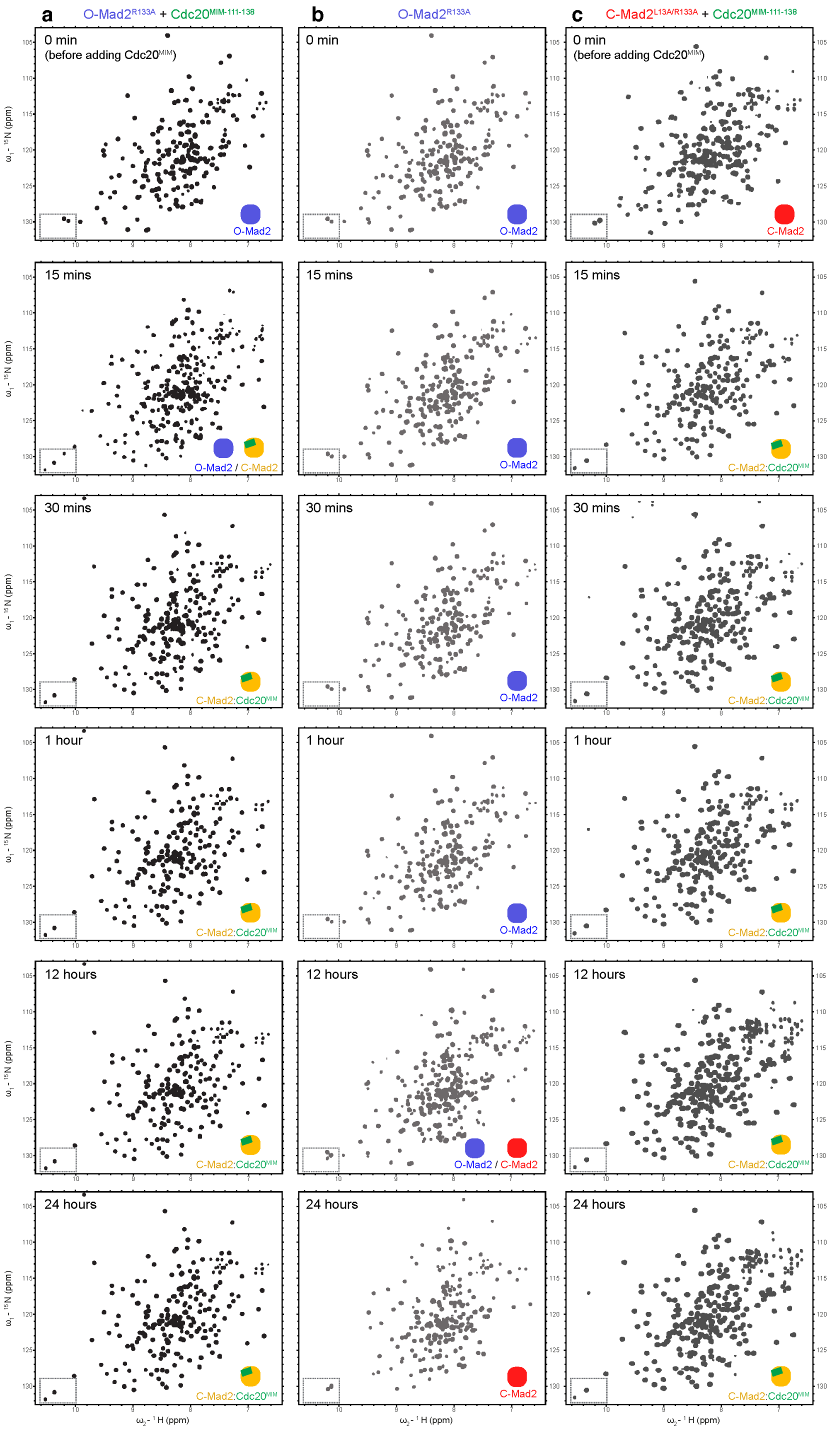


**Supplementary Figure 4. Conversion of O-Mad2 and 'empty' C-Mad2 to 'bound' C-Mad2.** **a.** Time-resolved HSQC time-course of O-Mad2$^{R133A}$ in the presence of two-fold molar excess of Cdc20$^{MIM\text{-}111\text{-}138}$ peptide. Conversion of Mad2 was traced over a 24 h time-course at 25 ˚C. **b.** HSQC time-course of O-Mad2 in the absence of MIM peptide. **c.** HSQC time-course of 'empty' C-Mad2$^{L13A/R133A}$ in the presence of two-fold molar excess of Cdc20$^{MIM\text{-}111\text{-}138}$ peptide. Side chains of Trp75 and Trp167 are diagnostic of Mad2 conformation, and their resonances are boxed on each spectrum. The schematics show the conformation of Mad2 as indicated by the tryptophan side-chain resonances. Data in **a** and **b** are reproduced from Supp. Fig. 13 of [31] for comparison.

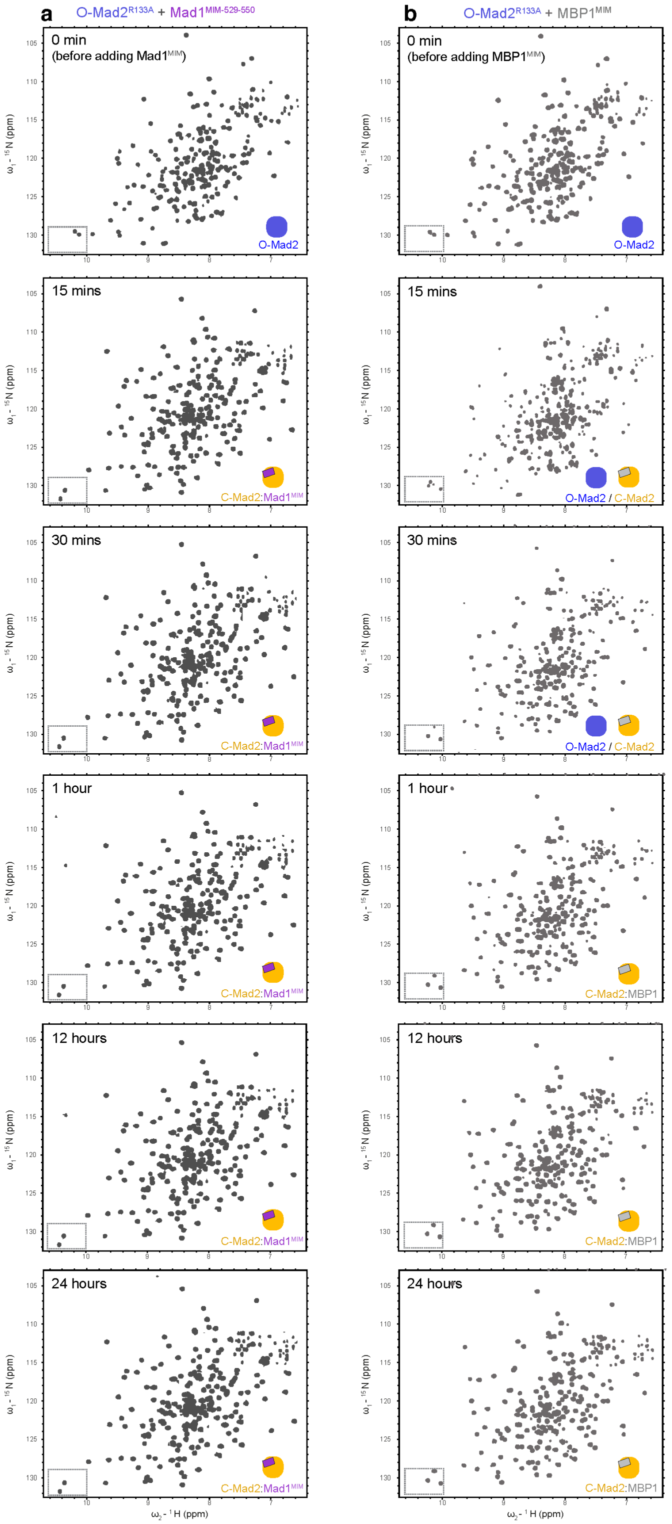


**Supplementary Figure 5. Mad2 conversion in the presence of different ligands. a.** Time-resolved HSQC time-course of O-Mad2$^{R133A}$ in the presence of two-fold molar excess of Mad1$^{MIM-529-550}$. Mad2 conversion was traced over a 24 h time course at 25 ˚C. **b.** HSQC time-course of O-Mad2$^{R133A}$ in the presence of two-fold molar excess of MBP1$^{MIM}$. Side chains of Trp75 and Trp167 are diagnostic of Mad2 conformation, and their resonances are boxed on each spectrum. The schematics show the conformation of Mad2 as indicated by the tryptophan side-chain resonances.

### Supplementary Figure 6

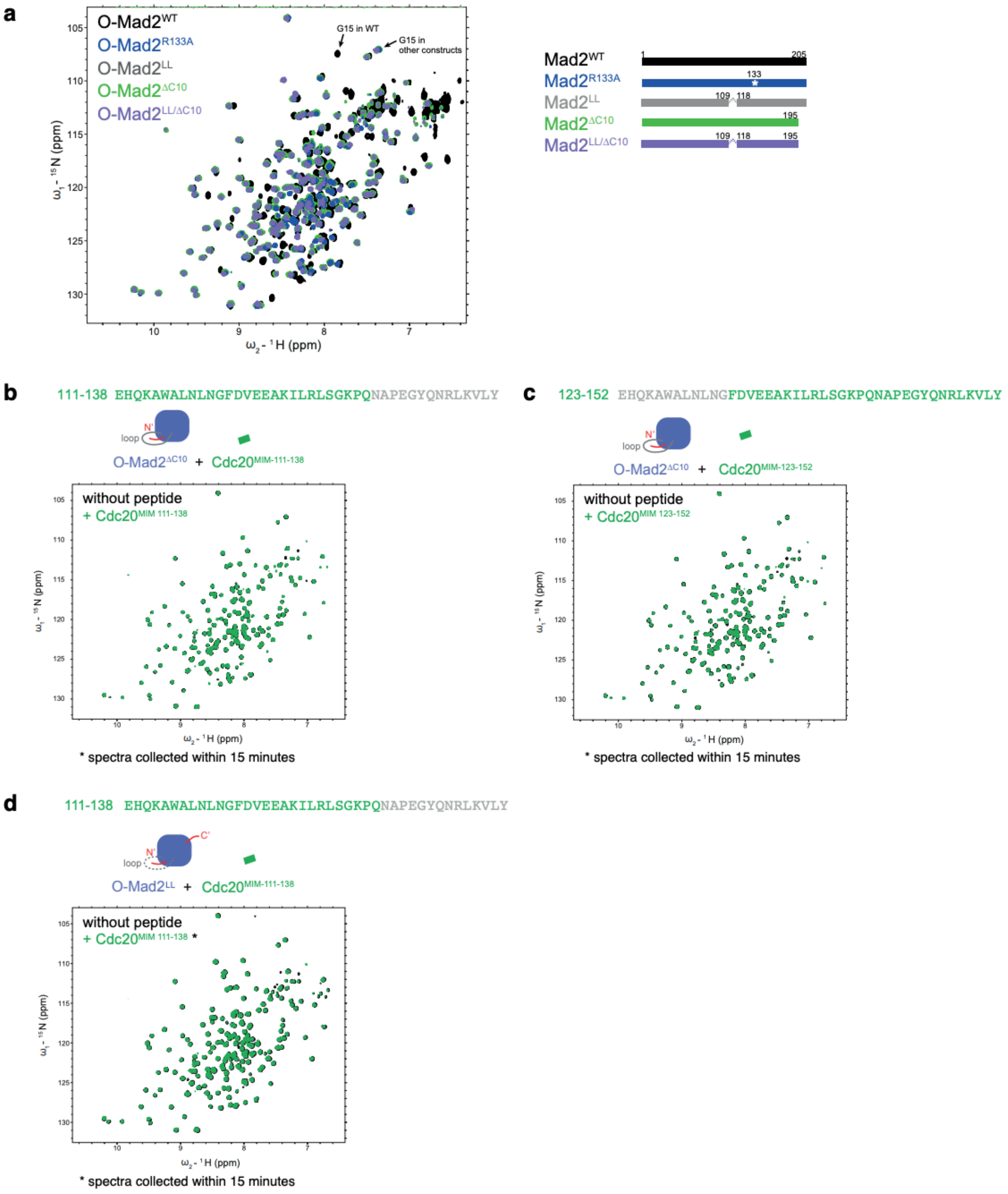


**Supplementary Figure 6. HSQC spectra of Mad2 variants and interaction between O-Mad2$^{\Delta C10}$ and Mad2$^{LL}$ with Cdc20$^{MIM}$. a.** HSQC of Mad2$^{WT}$ (black), Mad2$^{R133A}$ (blue) [31], Mad2$^{LL}$ (grey), Mad2$^{\Delta C10}$ (green) and Mad2$^{LL/\Delta C10}$ (purple). These variants were used to assign the backbone resonances of full-length O-Mad2. LL stands for loopless where residue 109-117 of the β5-αC loop were deleted, and ΔC10 lacks the C-terminal ten residues 196-205. Mad2$^{LL}$, Mad2$^{\Delta C10}$ and Mad2$^{LL+\Delta C10}$ were individually assigned, and their assignments were transferred to full-length Mad2$^{R133A}$. **b.** HSQC of O-Mad2$^{\Delta C10}$ in the absence (black) or presence (green) of a two-fold molar excess of Cdc20$^{MIM-111-138}$ peptide. **c.** HSQC of O-Mad2$^{\Delta C10}$ in the absence (black) or presence (green) of a two-fold molar excess of Cdc20$^{MIM-123-152}$ peptide. **d.** HSQC of O-Mad2$^{LL}$ in the absence (black) or presence (green) of a two-fold molar excess of Cdc20$^{MIM-111-138}$ peptide. Spectra were collected within 15 min, as Mad2 conversion was observed with longer incubation times (Fig. 5). No binding of Cdc20$^{MIM}$ to O-Mad2$^{\Delta C10}$ was observed indicating that the MIM-binding site is inaccessible in O-Mad2.

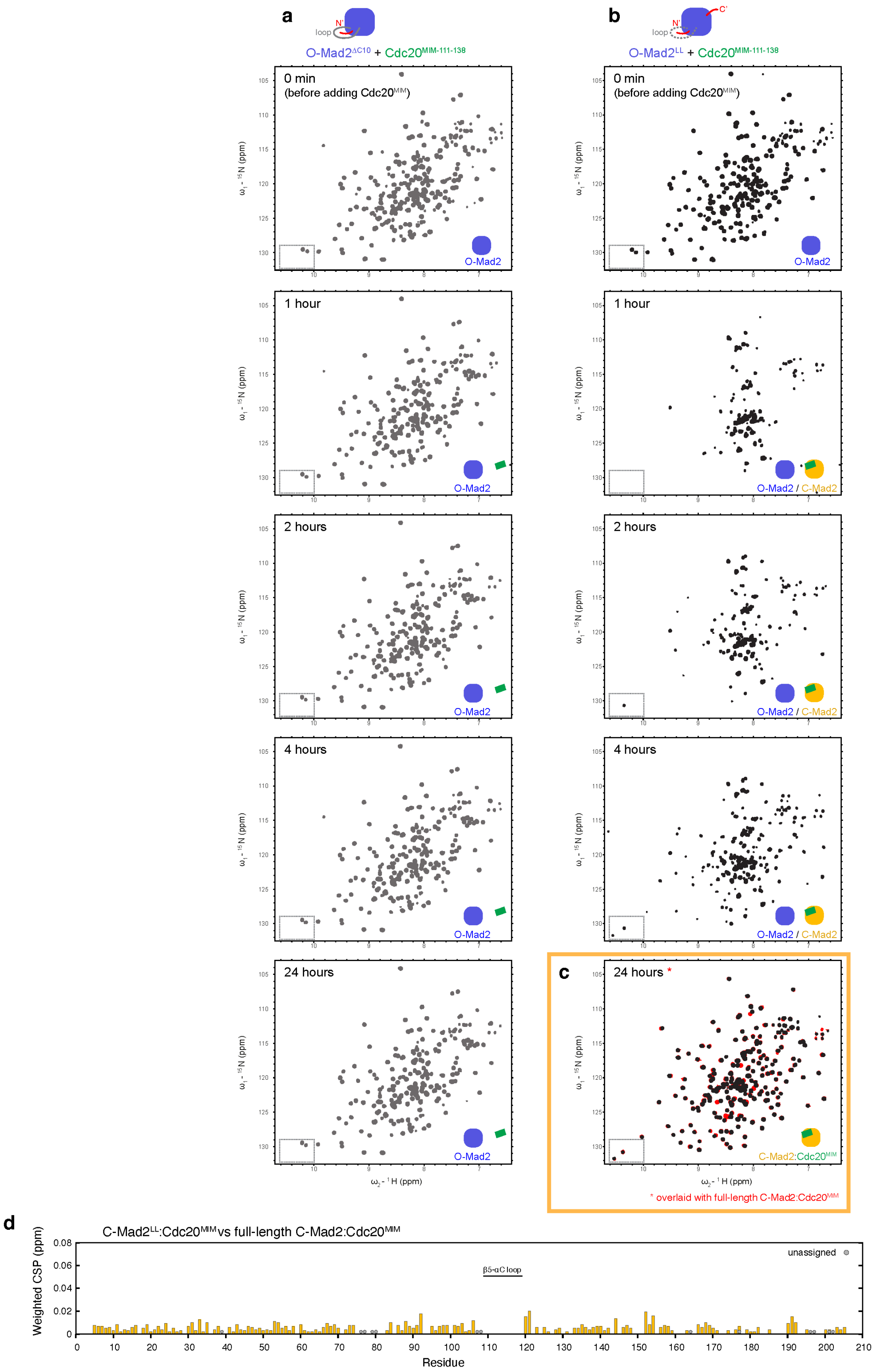


**Supplementary Figure 7. Conversion of O-Mad2 variants lacking the ten C-terminal residues (O-Mad2$^{\Delta C10}$) and β5-αC loop (O-Mad2$^{LL}$).** Mad2 conversion was traced over a 24-h time-course at 25 ˚C in the presence of two-fold molar excess of Cdc20$^{MIM\text{-}111\text{-}138}$ peptide. The schematics show the conformation of Mad2 as indicated by the tryptophan side-chain resonances. **a.** HSQC time-course of O-Mad2$^{\Delta C10}$. **b.** HSQC time-course of O-Mad2$^{LL}$. There was severe line broadening during conversion, mostly likely due to dimerisation. As this variant lacks the R133A mutation, O-Mad2$^{LL}$ would dimerise with the newly converted C-Mad2$^{LL}$. **c.** The final spectrum of O-Mad2$^{LL}$ + Cdc20$^{MIM-111-138}$ after 24 h (black) was overlaid with the spectrum of full-length O-Mad2$^{R133A}$ + Cdc20$^{MIM-111-138}$ after 24 h (red). Both variants converted to a 'bound' C-Mad2. **d.** The difference between the two spectra in **c** was analysed by their weighted CSPs. No significant CSPs were observed.

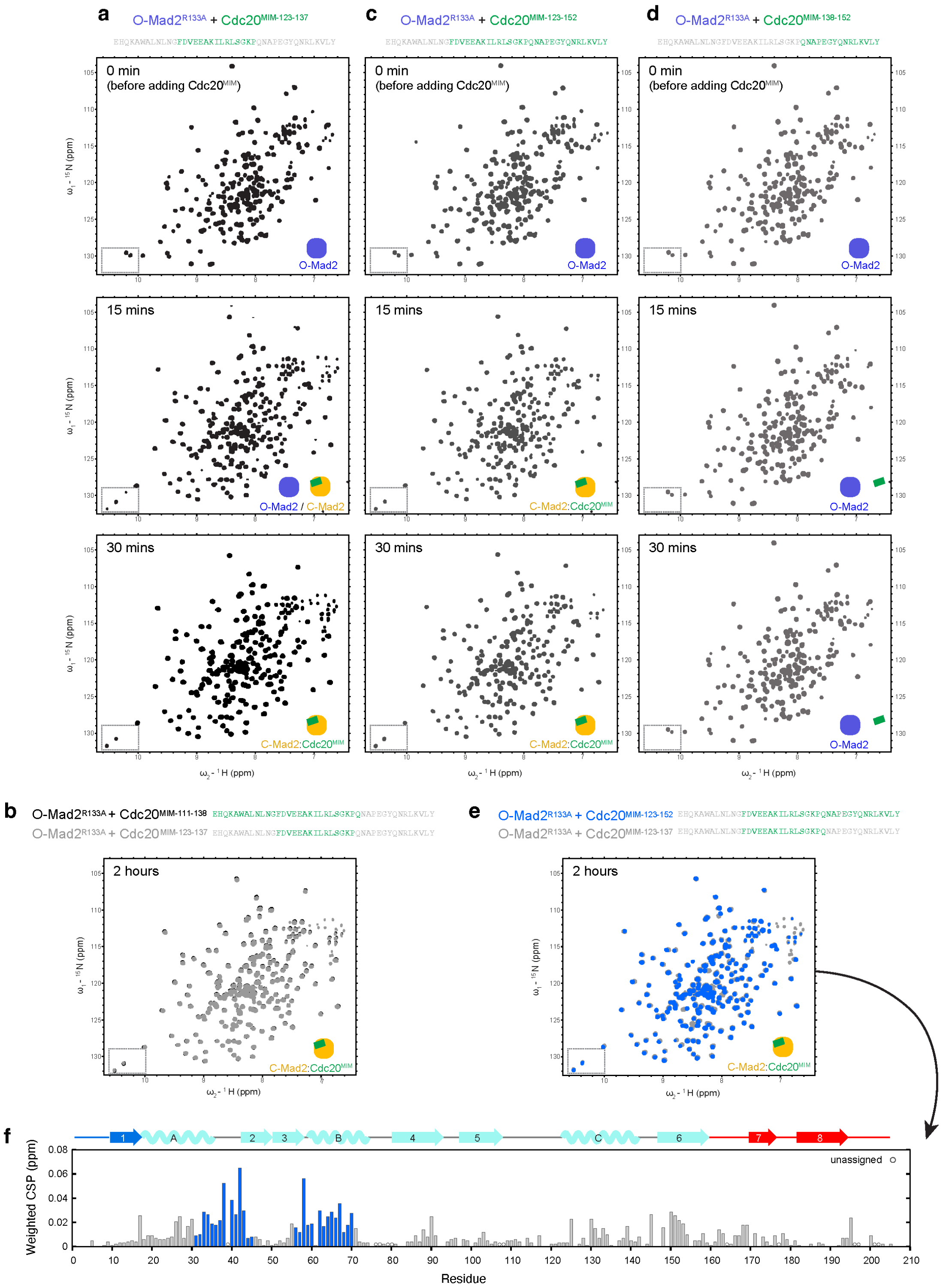

**Supplementary Figure 8. Mad2 conversion in the presence of various-length Cdc20$^{MIM}$ peptides.** **a.** HSQC time-course of O-Mad2$^{R133A}$ in the presence of two-fold molar excess of Cdc20$^{MIM-123-137}$. **b.** Overlay of the HSQC of O-Mad2$^{R133A}$ + Cdc20$^{MIM-111-138}$ (black) [31] and O-Mad2$^{R133A}$ + Cdc20$^{MIM-123-137}$ (grey) after 2 h. No significant changes were observed, suggesting Cdc20 residues 111-122 did not contribute to Mad2 binding. **c.** HSQC time-course of O-Mad2$^{R133A}$ in the presence of 2-molar excess of Cdc20$^{MIM-123-152}$. **d.** HSQC time-course of O-Mad2$^{R133A}$ in the presence of two-fold molar excess of

Cdc20$^{MIM\text{-}138\text{-}152}$. Mad2 conversion was traced over a 2 h-time course at 25 ˚C. Side chains of Trp75 and Trp167 are diagnostic of Mad2 conformation, and their resonances are boxed on each spectrum. The schematics show the conformation of Mad2 as indicated by the tryptophan side-chain resonances. The sequence for Cdc20 residues 111-152 is shown above the spectra, with the residues present in the peptide coloured green. **e.** Overlay of the HSQC of O-Mad2$^{R133A}$ + Cdc20$^{MIM\text{-}123\text{-}137}$ (grey) and O-Mad2$^{R133A}$ + Cdc20$^{MIM\text{-}123\text{-}152}$ (blue) after 2 h. The difference in the spectra indicate that Cdc20 residues 138-152 and residues 123-137 bind to different sites on O-Mad2. **f.** Analysis of the weighted CSPs observed in **e.** Unassigned residues are denoted by open circles. Residues involved in binding Cdc20$^{MIM\text{-}138\text{-}152}$ are highlighted in blue.

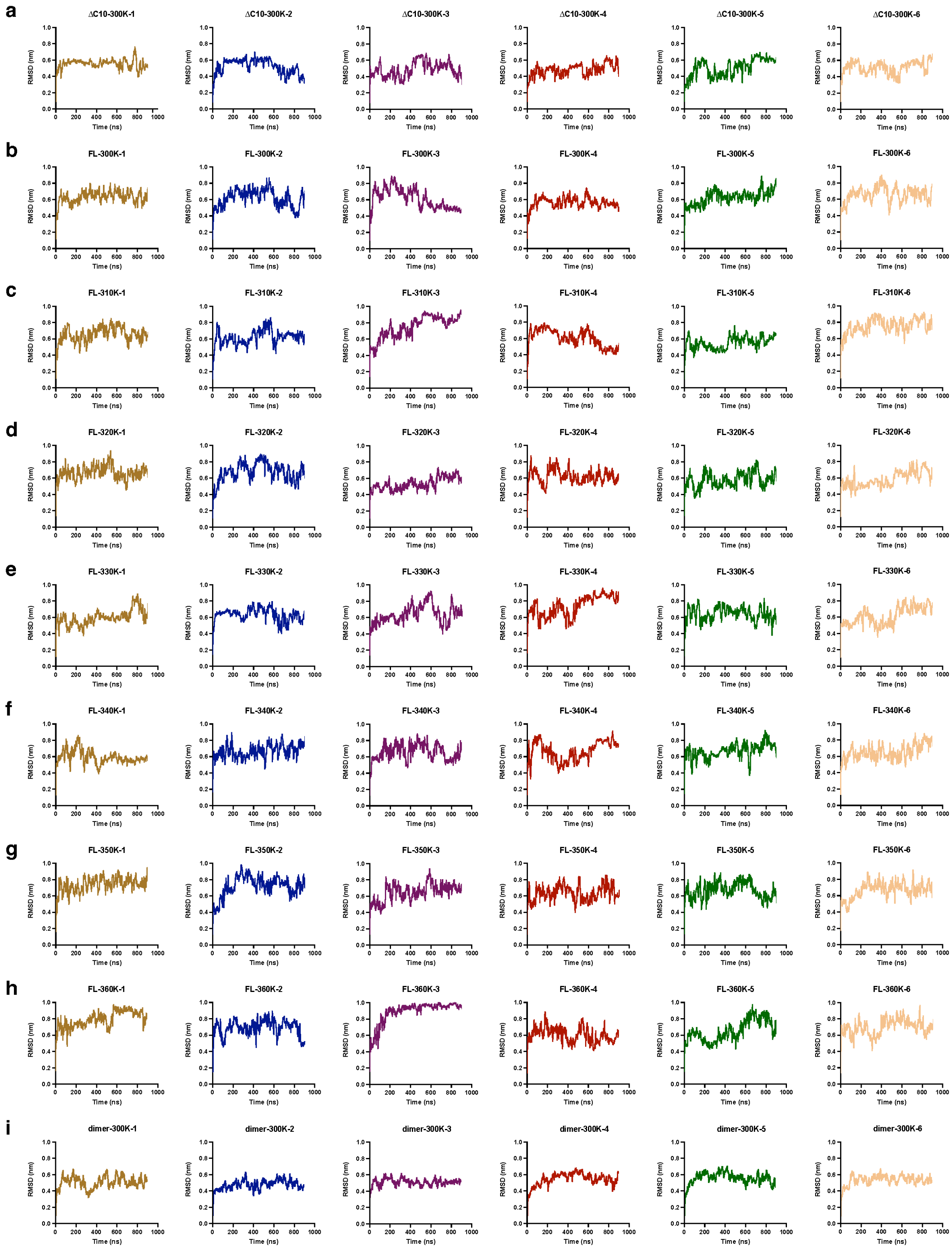


**Supplementary Figure 9. RMSD plots for MD simulations of O-Mad2.** Plots showing RMSD to the native structure for MD simulations of the following Mad2 variants: **a.** O-Mad2$^{\Delta C10}$ at 300 K, **b.** full-length O-Mad2 (FL) at 300 K, **c.** O-Mad2$^{FL}$ at 310 K, **d.** O-Mad2$^{FL}$ at 320 K, **e.** O-Mad2$^{FL}$ at 330 K, **f.** O-Mad2$^{FL}$ at 340 K, **g.** O-Mad2$^{FL}$ at 350 K, **h.** O-Mad2$^{FL}$ at 360 K and **i.** O-Mad2$^{FL}$:C-Mad2$^{FL}$:Mad1$^{MIM}$

(dimer) at 300 K. The starting structures for MD simulations on monomeric O-Mad2 were built using the crystal structure of O-Mad2$^{LL}$ (PDB 2V64, chain D) [16] and the missing segments (residues 1-8, 109-118, 195-205) were built using Modeller. The C-terminal 10 residues were manually truncated for the simulation of Mad2$^{\Delta C10}$. The starting structure for MD simulations on full-length O-Mad2:C-Mad2:Mad1$^{MIM}$ were built using the crystal structure of O-Mad2$^{LL}$:C-Mad2:MBP1$^{MIM}$ (PDB 2V64) [16]. The missing segments in O-Mad2 were built as previously described and MBP1$^{MIM}$ was manually modified to match the sequence of Mad1$^{MIM\text{-}538\text{-}551}$, guided by the structure of C-Mad2$^{R133A}$:Mad1$^{485\text{-}718}$ (PDB 1GO4). Cys79 and Cys106 were defined as unoxidised. Simulations were carried out using the DES-Amber force field and TIP4P-D water model. A total of 5.4 µs of simulations were carried out for each variant at each temperature, using six independent simulations of 0.9 µs. Cα RMSF was calculated from each trajectory and the average Cα RMSF is shown in Figs. 4 and 6e and Supplementary Fig. 10f.

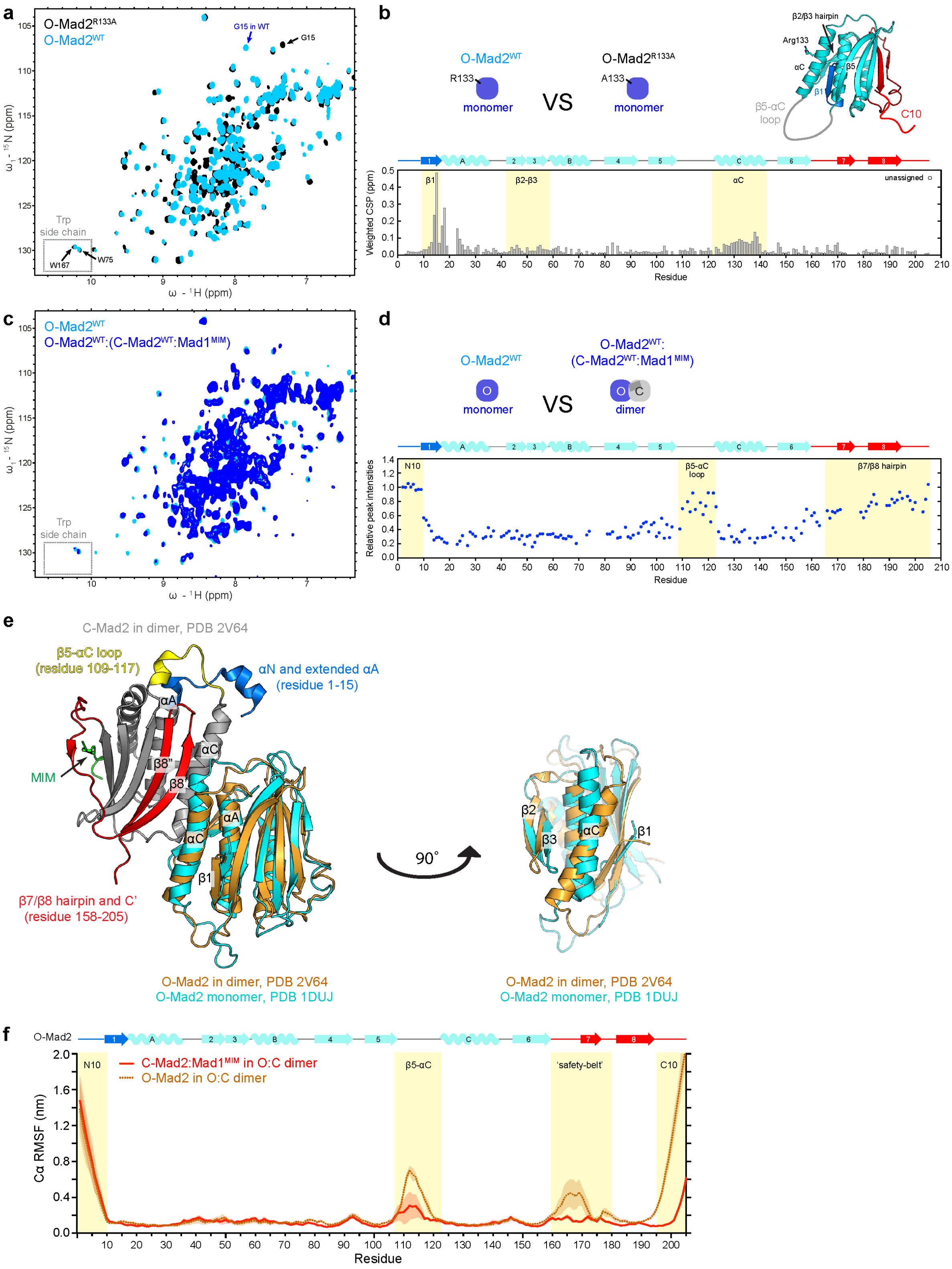


**Supplementary Figure 10. Comparison of O-Mad2 in different oligomeric states.** **a.** Overlay of HSQC of O-Mad2$^{R133A}$ (black) [31] and O-Mad2$^{WT}$ (light blue). Significant CSPs were observed in the β1 strand, as demonstrated by the resonances of Gly15. **b.** Analysis of the weighted CSPs observed in **a.**

Unassigned residues are denoted by open circles. Regions of interest are indicated in the highlighted residues. Model of the full-length O-Mad2 is shown to indicate the position of Arg133. **c.** Overlay of HSQC of monomeric O-Mad2$^{WT}$ (light blue) and dimeric O-Mad2$^{WT}$ (blue). Dimeric O-Mad2$^{WT}$ was prepared by mixing isotopically labelled O-Mad2$^{WT}$ and excess unlabelled C-Mad2$^{WT}$ bound to Mad1$^{MIM}$. **d.** Relative peak intensities in **c.** were analysed and mapped onto the Mad2 sequence. Peak intensities were normalised to the C-terminal residue Asp205. Significant line broadening was observed across the sequence upon dimerisation except for the N-terminal ten residues (N10), β5-αC loop and the β7/β8 hairpin, which are highlighted in yellow. **e.** The structure of O-Mad2$^{LL}$:C-Mad2:MBP1$^{MIM}$ (PDB 2V64) [16] is shown with O-Mad2$^{LL}$ (brown) overlaying the solution structure of monomeric O-Mad2$^{\Delta N10}$ (cyan, PDB 1DUJ) [12]. The C-terminal residues 196-205 are absent in the solution structure. C-Mad2 is coloured in grey and the MIM is coloured in green. Upon the O- to C-Mad2 conversion, the C-terminal residues 196-200 become part of the β8" strand and the newly formed 'safety-belt' wraps around the protein core (red), resulting in a decrease in their dynamics. The flexibility of the β5-αC loop (yellow) is also restrained by the extended αA helix (blue). The N-terminal ten residues are shown here to adopt a short αN helix (blue), but likely it remains unstructured and highly dynamic in solution (Supplementary Figs. 3d). (Right) The O-Mad2 structures are also viewed from a different angle to highlight the difference in their αC helix and β2/β3 hairpin. The αC helix and β2/β3 hairpin of O-Mad2 shift position to avoid a steric clash with αC of C-Mad2. These rearrangements likely facilitate the release of β1 strand during O- to C-Mad2 conversion. **f.** Cα RMSF were determined from MD simulations at 300 K for O-Mad2 (brown dotted line) and C-Mad2:Mad1$^{MIM}$ (red solid line) in a O-Mad2:C-Mad2:Mad1$^{MIM}$ dimer. A total of 5.4 μs of simulations were carried out for each variant, using six independent simulations of 0.9 μs. RMSD plots for all MD simulations are shown in Supplementary Fig. 10. The continuous error bands indicate one standard deviation derived from six independent simulations. O-Mad2 shows distinct dynamic features that are absent in C-Mad2, including a highly dynamic β5-αC loop, residues 160-180 that constitute the 'safety-belt' in C-Mad2, and the C-terminal ten residues. However, the N-terminal ten residues remain highly dynamic, in agreement with the absence of helical propensity observed in C-Mad2:Cdc20$^{MIM}$ (Supplementary Fig. 3d).

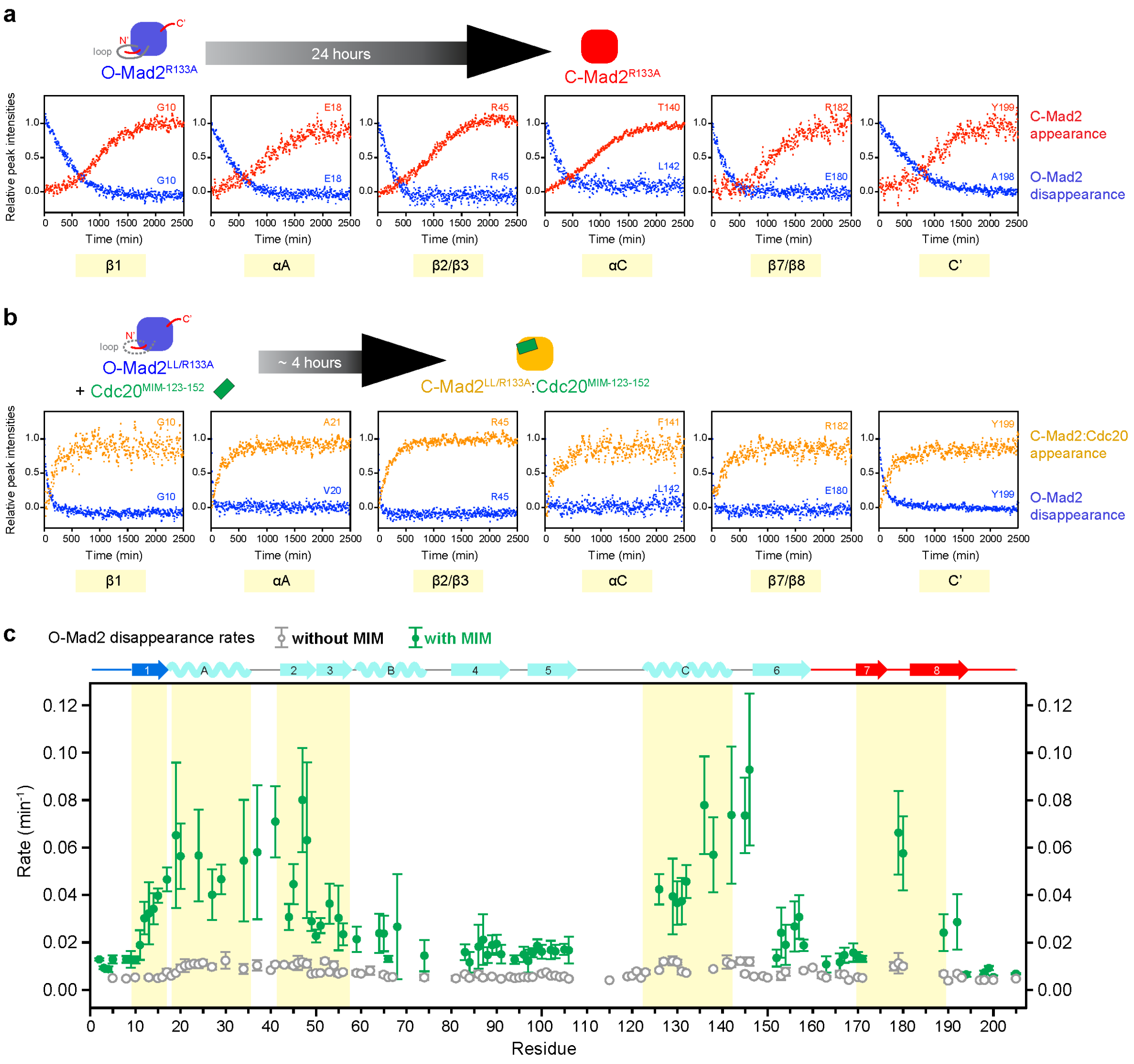

**Supplementary Figure 11. Tracing Mad2 conversion by NMR. a, b.** Consecutive 2D NMR spectra were collected at 25 ˚C across 42 h to trace the conversion of full-length O-Mad2$^{R133A}$ in the absence of Cdc20$^{MIM}$ (**a**) and an O-Mad2$^{LL/R133A}$ mutant that has a reduced conversion rate of ~250 mins in the presence of Cdc20$^{MIM-123-152}$ (**b**). The blue dots are the relative peak intensities of O-Mad2 resonances. The red dots in **a** and orange dots in **b** are the relative peak intensities of 'empty' C-Mad2 resonances and C-Mad2:Cdc20$^{MIM}$ resonances respectively. The changes in peak intensities for representative residues from the N-terminal β1 strand, αA helix, β2/β3 hairpin, αC helix, β7/β8 hairpin and the C terminus are shown. When available, data from the same residue of O-Mad2 and C-Mad2 are shown for comparison (e.g. G10 in β1). Otherwise, data from a neighbouring residue are shown (e.g. V20 and A21 in αA). In all residues, the C-Mad2 resonances appeared at a lower rate than the disappearance of O-Mad2 resonances, suggesting there are intermediate states during Mad2 conversion. **c.** The initial conversion rates of individual O-Mad2 residues were determined by plotting the peak intensities of their O-Mad2 resonances against time and fitting the data to an exponential decay curve. Error bars represent the standard regression errors of the fit. The rates for O-Mad2$^{R133A}$ in the absence of MIM were plotted as grey open circles (without MIM) and those for O-Mad2$^{LL/R133A}$ in the presence of Cdc20$^{MIM\ 123-152}$ were plotted as green circles (with MIM). The data were scaled for comparison between the varying conversion rates across the sequence in Fig. 7d.
